# Substrate recognition principles for the PP2A-B55 protein phosphatase

**DOI:** 10.1101/2024.02.10.579793

**Authors:** Thomas Kruse, Dimitriya H. Garvanska, Julia Varga, William Garland, Brennan McEwan, Jamin B. Hein, Melanie Bianca Weisser, Iker Benavides Puy, Camilla Bachman Chan, Paula Sotelo Parrila, Blanca Lopez Mendez, Jeyaprakash Arulanandam, Ora Schueler-Furman, Torben Heick Jensen, Arminja Kettenbach, Jakob Nilsson

## Abstract

The PP2A-B55 phosphatase regulates a plethora of signaling pathways throughout eukaryotes. How PP2A-B55 selects its substrates presents a severe knowledge gap. By integrating AlphaFold modelling with comprehensive high resolution mutational scanning, we show that α-helices in substrates bind B55 through an evolutionary conserved mechanism. Despite a large diversity in sequence and composition, these α-helices share key amino acid determinants that engage discrete hydrophobic and electrostatic patches. Using deep learning protein design, we generate a specific and potent competitive peptide inhibitor of PP2A-B55 substrate interactions. With this inhibitor, we uncover that PP2A-B55 regulates the nuclear exosome targeting complex by binding to an α-helical recruitment module in RBM7. Collectively, our findings provide a framework for the understanding and interrogation of PP2A-B55 in health and disease.

**One sentence summary:** α-helices in PP2A-B55 substrates bind a conserved pocket on B55 through a common mechanism that is conserved in eukaryotes.

## Introduction

Dynamic phosphorylation of Ser/Thr residues is central for modulating protein activities throughout eukaryotes (*1, 2*). This regulatory mechanism depends on the selective recognition of substrates and substrate residues by kinase and phosphatase enzymes. While the basis for kinase specificity has been extensively characterized (*3, 4*), the parameters mediating protein phosphatase specificity are less well understood. In recent years it has become clear that members of the Ser/Thr phosphoprotein phosphatases (PPPs; PP1-PP7) achieve specificity by binding to short linear motifs (SLiMs) in the intrinsically disordered regions of their substrates or substrate specifiers (*5–7*). SLiMs bind to defined pockets distal from the active site on the phosphatase catalytic subunits. However, whether SLiMs mediate substrate recognition by all PPP members is not known.

The phosphoprotein phosphatase PP2A regulates numerous fundamental signaling pathways and is deregulated in human diseases underscoring the importance of establishing fundamental principles of PP2A substrate recognition (*8–10*). PP2A holoenzymes are trimeric and composed of a catalytic subunit (PPP2CA/B), a scaffolding subunit (PPP2R1A/B) and one of several regulatory B subunits present in multiple isoforms (B55α-d, B56α-e, PR72/PR130, STRN1-4) (*6, 7, 11*). The B55 and B56 regulatory subunits are fully conserved from yeast to human, acting as major determinants of PP2A holoenzyme specificity. We and others previously uncovered a conserved binding motif for PP2A-B56, the so-called LxxIxE motif, that engages a highly conserved binding pocket on the B56 regulatory subunit (*12–14*). This discovery provided important insight into PP2A-B56 regulation of signaling and opened up new possibilities for the precise engineering of signaling pathways.

In contrast to PP2A-B56, we have a limited understanding of PP2A-B55 substrate recognition. PP2A-B55 is a key regulator of multiple cellular processes such as the cell cycle where it antagonizes cyclin dependent kinase activity. In turn, PP2A-B55 activity is tightly regulated by the ENSA/ARPP19 and FAM122A proteins, ensuring proper cell cycle execution (*15–19*). Recently, the p107 protein was proposed to bind B55 via a SLiM (*20*), while cryo-EM structures of the B55 inhibitors ARPP19 and FAM122A revealed binding of α-helices to B55 (*21*). It is unclear whether substrates bind to B55 via SLiMs, while inhibitors use a distinct mechanism.

Here we have combined AlphaFold modeling with experimental validation, which identified B55-interacting α-helices in numerous proteins conferring substrate specificity to PP2A-B55. Our discoveries have important implications for understanding and engineering PP2A signaling networks and provide a first example of an extended structured binding element conferring specificity to a PPP family member.

## Results

### A conserved pocket on the B55 subunit binds α helices in interactors

Despite the considerable amount of structural and biochemical information available on PP2A-B55, a key unresolved question is the molecular basis for its substrate recognition. To address this question, we established a discovery pipeline that integrates AlphaFold2 (AF) multimer modelling with experimental high-resolution mutagenesis scanning of PP2A-B55 interacting proteins (Fig. 1A). AF can not only generate highly accurate models, but also provides confident measures of the model quality (*22–25*). As input for the pipeline we used our previous mass spectrometry interactome data that identified 256 protein interactors specific for PP2A-B55 (*14, 26*). We expanded this with reported B55 binding domains from the literature (EYA3, AMOTL2) as well as instances from yeast (Zds1) and viruses (E4ORF4) (*27–31*). Applying AF on this set of interactors returned 40 models with good confidence scores (iPAE< 10 and average interface pLDDT > 70, except EYA3 and AMOTL2 that was included because of previous validation of binding domains, Fig. S1A, Supplemental Table 1 and Supplemental Dataset 1). The majority of models revealed defined binding elements predicted to engage a fully conserved surface on B55 that has previously been implicated in substrate binding (*19, 20, 32*). The recently reported binding elements in FAM122A and ARPP19/ENSA were also identified through this approach (*21*). The AF models for FAM122A and ARPP19 overlap at the atomic level with the experimental structure for FAM122A, as well as for the helix covering the L49/L53 motif in ARPP19, providing confidence in our AF models (Fig. S1B-C).

**Figure 1.**
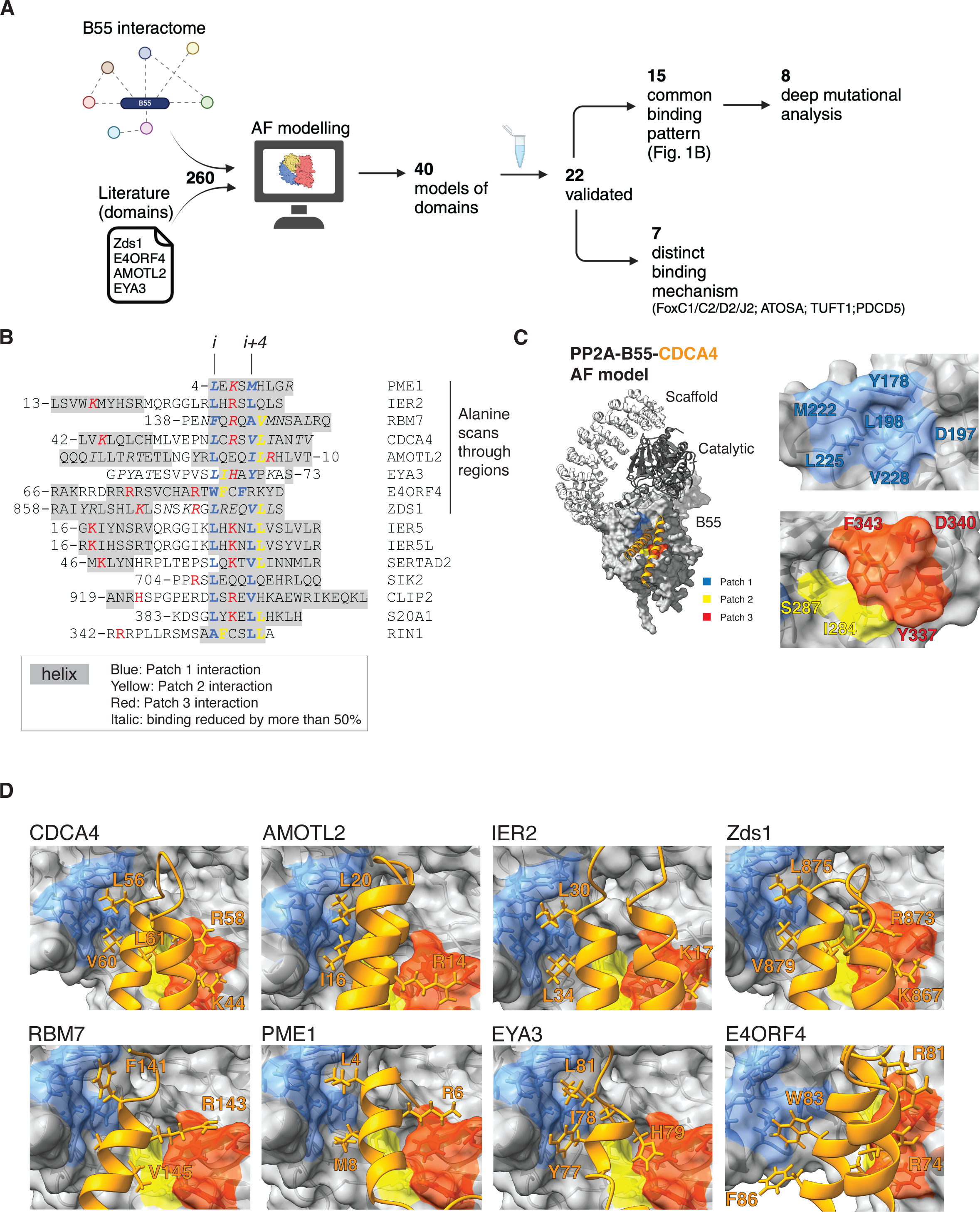
Helical motifs engage a conserved binding pocket on PP2A-B55. **A**) Schematic of pipeline to identify PP2A-B55 binding elements. **B**) Alignment of validated instances with helices in grey, residues contacting patch 1 in blue, patch 2 in yellow and patch 3 in red. The first 8 proteins on the list covers the proteins that was scanned by single Alanine mutagenesis and residues in italic was found to reduce binding by at least 50%. **C**) Model of the PP2A-B55-CDCA4 complex and the different patches in B55 indicated. **D)** AF models of the indicated proteins and their interaction with B55. Core residues interaction with the different patches in stick.

Out of the predicted 40 binding elements we experimentally validated 19 instances, using immunopurifications from HeLa cells (Fig. S2-S10). To this list we added 3 additional instances based on high sequence homology (SERTA domain of SERTAD2 and IER5/IER5L), collectively expanding the validated and modeled B55 binding elements to 22.

We inspected the 22 AF models for common features, which revealed that all of them used α-helical elements to bind B55. The presence of instances in yeast, humans, and viruses suggests an evolutionary conserved binding mechanism. Consistently, SERTA domains from numerous species spanning close to a billion years of evolution were observed to bind PP2A-B55 (Fig. S11). For 15 binding elements, we identified a common pattern of binding with either a single helix (PME1, RBM7, EYA3, SIK2, S20A1, RIN1) or helix-loop-helix (CDCA4, SERTAD2, IER2/5/5L, Zds1, CLIP2, AMOTL2) engaging B55 (Fig. 1B-D). E4ORF4 also used this binding mechanism through a three-helix bundle. In all cases, a single helix, the binding helix, binds to B55 in a similar manner. In the case of helix-loop-helix binders and E4ORF4 additional contacts with B55 are established via the preceding additional helix (or helices, in the case of E4ORF4) (Fig.S10).

Despite limited sequence similarity, our structural analysis uncovered general principles for how the binding helix engages B55. These principles involve a conserved contact of a central motif at the start of the binding helix with an adjacent hydrophobic pocket composed of B55 residues Y178, D197, M222, L225 and V228) that we term *patch 1* (Fig. 1B-D). This central motif involves primarily hydrophobic residues at positions *i* and *i+4* in the helix (e.g. L56/V60 in CDCA4). This patch is complemented by *patch 2,* composed of B55 residues I284 and S287, which is contacted by residues at position i+1 and/or i+5 that can be hydrophobic (such as L61 in CDCA4A). *Patch 3* is centered around B55 residue D340 that forms electrostatic interactions predominantly with the residue located on the opposite side of the binding helix, at position *i+*2 (e.g. R58 in CDCA4). In the helix-loop-helix type of binders, the *i+*2 position is supplemented by additional positively charged residues (e.g. K44 in the adjacent helix of CDCA4). The interactions with *patch 3* display more variation, and may involve residues contributed from different parts of the substrate, e.g. in Zds1 where residue *i+2* (E27) is negatively charged and R23 in a nearby loop forms the salt bridge with D340 instead. Besides these two patches, additional contacts are formed with the side of the binding pocket involving aromatic B55 residues Y337 and F343. Although our AF models support these general principles of binding, they also revealed distinct additional contacts to B55 for each binder. Seven of our 22 validated instances (FOXC1/C2/D2/J2, ATOSA, TUFT1 and PDCD5) also engaged B55 through α-helical structures but in a more heterogenous manner not readily conforming to the common binding patterns observed above. However, the AF models revealed that they still contacted *patch* 1-3, underscoring the importance of these patches for binding (Fig. S2).

To obtain experimental support for the AF models, we conducted comprehensive biochemical mappings involving truncation analysis, and ten and/or five alanine walks through several interactors mapping regions of interaction consistent with the AF models (Fig. S3-5, S7-9). To further identify specific amino acid binding determinants at the B55-interactor interface, single alanine mutational scans of these interfaces were performed for CDCA4, IER2, RBM7, AMOTL2, EYA3, PME1, Zds1 and E4ORF4 (Fig. 1B, Fig. S3-10). For the majority of instances, the contribution of key amino acid residues interacting with the different patches of B55 as modelled by AF was confirmed (Fig. 1B). Through mass spectrometry analysis we confirmed that the binding of several isoforms of B55 was abolished when we mutated key binding residues in CDCA4, IER2, PME1, AMOTL2 and RBM7 in line with the conservation of the B55 binding pocket (Fig. S12, Supplemental Table 2).

To investigate the contribution of the different B55 patches, we analysed the binding of PME1, FAM122A, IER2, and CDCA4 to a panel of B55 mutants covering the different patch residues (Fig. 2A-B, Fig. S13). This revealed substantial diversity of these proteins in their binding requirements for the three patches. PME1 required amino acids in all three patches for binding, whereas only residues in patch 1 seemed necessary for FAM122A. IER2 and CDCA4 shared similar binding characteristics with requirements from residues in patch 1 and 3. To explore whether this patch variability is a general phenomenon shared with other B55 interactors we performed mass spectrometry analysis of B55 WT, V228A, I284A, S287A and D340A immunopurifications (Figure 2B). The results confirmed on a global scale that B55 interactors have differences in dependencies for the different patches on B55, despite common features of the binding mechanism (Figure 2C and supplemental Figure S13). This is exemplified by IER2 versus IER5 binding to B55. Despite their almost identical N-terminal helix-loop-helix structure IER5 requires S287 in *patch 2* for efficient interaction whereas IER2 does not (Figure 2C).

**Figure 2.**
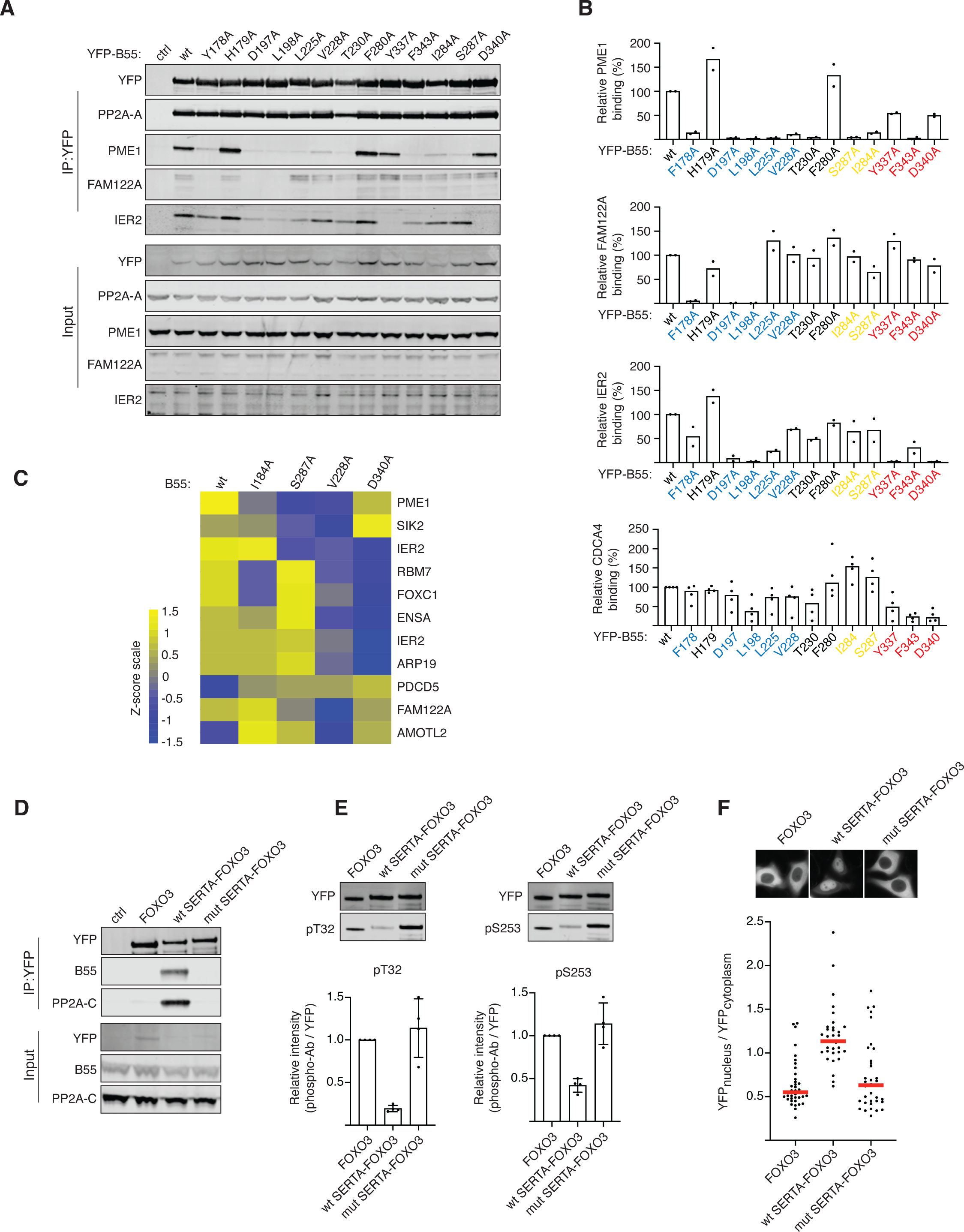
Helical motifs can act as substrate specifiying elements. **A**) The indicated B55 mutants was purified and binding to IER2, PME1, CDCA4 and FAM122A determined and quantified.B) Quantifications of relative binding of proteins indicated to B55. **C**) Heat map illustrating the changes in binding pattern of the indicated proteins to the different B55 variants as determined by mass spectrometry. The scale is log2. **D)** The CDCA4 wt or mutant SERTA domain was fused to FOXO3 and binding to PP2A-B55 monitored by western blot. **E)** As **D**) but IPs probed with FOXO3 phosphospecific antibodies. **F)** Localisation of FOXO3 fusion proteins by live cell microscopy.

In summary, our AF models and experimental data collectively support that α-helices in binders share key amino acid determinants that differentially engage discrete hydrophic and electrostatic patches on B55 making it a remarkably versatile platform for mediating protein-protein interactions.

### Helical binding elements confer substrate specificity to PP2A-B55

An important outstanding question was whether the α-helices acted as substrate specifiers or inhibitors of PP2A-B55. Our mass spectrometry analysis revealed that a number of phosphorylation sites in RBM7 (Fig.S5A), PME1 (Fig.S6A) and AMOTL2 (Fig.S8A) and had a higher degree of occupancy when we mutated the B55 binding helix, which would support a role in substrate specification (Supplemental Table 2). To directly test this, we engrafted the SERTA domain from CDCA4 or a mutant variant (SERTA K44A/V60A/L61A) onto the FOXO3 transcription factor on which we mutated its endogenous PP2A-B56 binding site (*14*) (FOXO3 2A). We previously showed that PP2A-B56 regulates FOXO3 through binding to its LxxIxE motif. When this motif is mutated, dephosphorylation of FOXO3 T32 and S253 and nuclear translocation are prevented. The engraftment of the SERTA WT domain onto FOXO3 2A allowed efficient recruitment of PP2A-B55. In contrast, no binding of PP2A-B55 to FOXO3 2A fused to the mutated SERTA domain could be detected (Fig. 2D). Strikingly, we observed efficient dephosphorylation of FOXO3 T32 and S253 when we engrafted SERTA WT, and this resulted in nuclear translocation of FOXO3 as predicted (Fig. 2D-F). We conclude that α-helical recruitment modules can serve as substrate specifying elements for PP2A-B55.

### Generation of an engineered B55 inhibitor blocking binding of helical elements

Our data revealed that the conserved surface on B55 engages numerous substrates. Based on this we predicted that a high affinity peptide binding to this surface on B55 would inhibit PP2A-B55 through competitive substrate displacement.

To generate this peptide, we turned to protein engineering using ProteinMPNN (*33*) and Rosetta FlexPep Design (*34*) to generate a panel of artificial designs predicted to bind to PP2A-B55 as helix-loop-helix motifs (Fig. 3A). A screen for binding to PP2A-B55 in cells by affinity purifying tagged versions of the peptides revealed that all peptides designed with ProteinMPNN bound, while only one of the FlexPep designs bound (Fig. 3A). The strongest binder stood out by the highest pLDDT and Protein MPNN scores, and also resulted in the smallest RMSD to the starting structure upon refolding with AF2 ((a measure used to estimate whether the design indeed will adapt the structure it was designed to adapt, Fig. S14). We moved forward with the two strongest binders and measured their B55 binding affinity by surface plasmon resonance (SPR), and compared this to corresponding B55i CTRLs, in which residues crucial for the interaction were mutated. This revealed extremely tight binding of the inhibitors. In particular, inhibitor 1, termed B55i, showed strong B55 binding with an estimated K_d_ of 100 pM due to a low k_off_ rate (Fig. 3B and Fig. S14D). This strong binding was supported by fluorescent polarisation assays that measured a K_d_ of 6 nM (Fig. S14E). The tight binding of B55i is achieved through several positively charged residues contacting negative patches on B55 and good packing of the hydrophobic core. Despite considerable efforts in both recombinant production and peptide synthesis, we have been unable to compare these affinity measurements to the helix-loop-helix binding elements identified in CDCA4, IER2 and AMOTL2. This is probably due to the highly insoluble nature of these protein elements. However, we did manage to produce several of the single helix-type of B55 binders by protein expression and peptide synthesis and measured binding of PME1, Arpp19 and RBM7 to B55 in the low micromolar affinity range using the same SPR approach as for B55i (Fig. S15). To test the specificity of B55i in cells, we affinity purified YFP-tagged versions of B55i and B55i CTRL and analysed samples by colloidal staining and mass spectrometry (Fig. 3C, Fig. S14F and Supplemental Table 2). This revealed a strong and very specific enrichment of all PP2A-B55 isoforms by B55i that was entirely abrogated by the mutations. These experiments emphasize the strong affinity and high selectivity of the designed inhibitor peptide for PP2A-B55 complexes.

**Figure 3.**
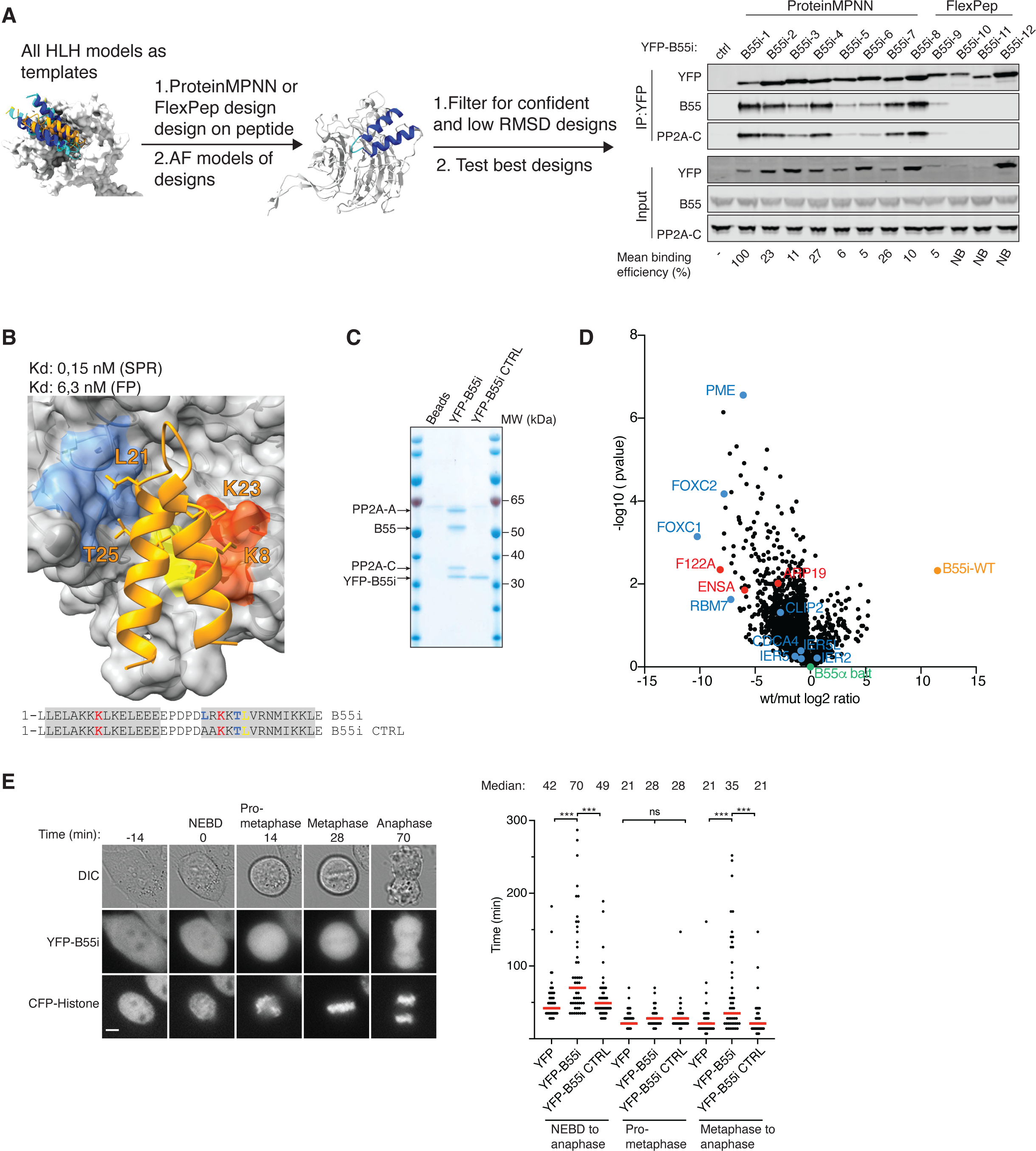
Generation of a specific B55 inhibitor. **A)** Design pipeline for generating a specific and tight binder of B55 and test of the top designs by immunopurification. **B**) Model of the B55i bound to B55 and measured Kd and Ki indicated below. **C**) Coomassie stained gel of B55i and B55i CTRL purified from HeLa cells. **D**) B55 was affinity purified after incubation with either B55i or B55i CTRL peeptides and samples analysed by mass spectrometry. **E**) Mitotic duration in cells expressing B55i measured by timelapse microscopy. Scale bar is 5 μM. Right panel: Quantification of **E**). Shown is pooled data from three independent experiments. Each circle represents the timing of a single cell. Red line indicates the median time. Mann–Whitney U-test was applied. ns, not significant. \*\*\**P* < 0.001.

To test the ability of the inhibitor to displace substrates from PP2A-B55, we affinity purified B55 from cells in the presence of either B55i or B55i CTRL peptides and analysed complexes by mass spectrometry. This revealed that B55i specifically bound the complex and reduced the binding of several interactors containing a B55 binding site as well as the endogenous B55 inhibitors Arpp19, ENSA and FAM122A (Fig. 3D, Supplemental Table 2). To test if B55i could block the biological function of PP2A-B55, we expressed YFP-tagged versions of B55i and B55i CTRL in HeLa cells and analysed their effect on mitotic exit, which is a cellular process that depends on PP2A-B55 activity (*35, 36*). Using live cell time-lapse microscopy we observed a delay in the metaphase to anaphase transition in cells expressing B55i inhibitor, while the duration of prometaphase was not affected (Fig. 3E). Thus, B55i causes *in vivo* phenotypes in line with the known function of PP2A-B55.

In conclusion, we used de novo protein design to generate a highly potent and specific PP2A-B55 inhibitor providing an important tool for dissecting and manipulating PP2A-B55 regulated signaling.

### PP2A-B55 regulates NEXT complex function by binding a helical domain of RBM7

Our characterization of PP2A-B55 binding elements allowed us to address new functions of this phosphatase. One of the helical binding elements that we mapped is located in the RBM7 protein that forms part of the nuclear exosome targeting (NEXT) complex. NEXT is composed of RBM7, ZCCHC8 and MTR4 (MTREX) and acts as an adaptor by channeling non-polyadenylated RNA to the ribonucleolytic RNA exosome complex for degradation (*37, 38*). The phosphorylation of RBM7 in DNA damage conditions has previously been shown to impair normal function of the NEXT complex in RNA decay (*39–41*), suggesting a regulatory control mechanisms mediated by phosphorylation/dephosphorylation. To explore a possible role of PP2A-B55 in regulating NEXT complex function, we induced the expression of B55i, or its mutant variant, in HeLa cells and monitored the levels of NEXT complex RNA substrates. This revealed a specific accumulation of the NEXT targets *proRBM39*, *proDNAJB4* and *proDDX6* when B55i, but not the control, was induced, supporting an impact of PP2A-B55 on NEXT complex function (Fig. 4A and Fig. S16).

**Figure 4.**
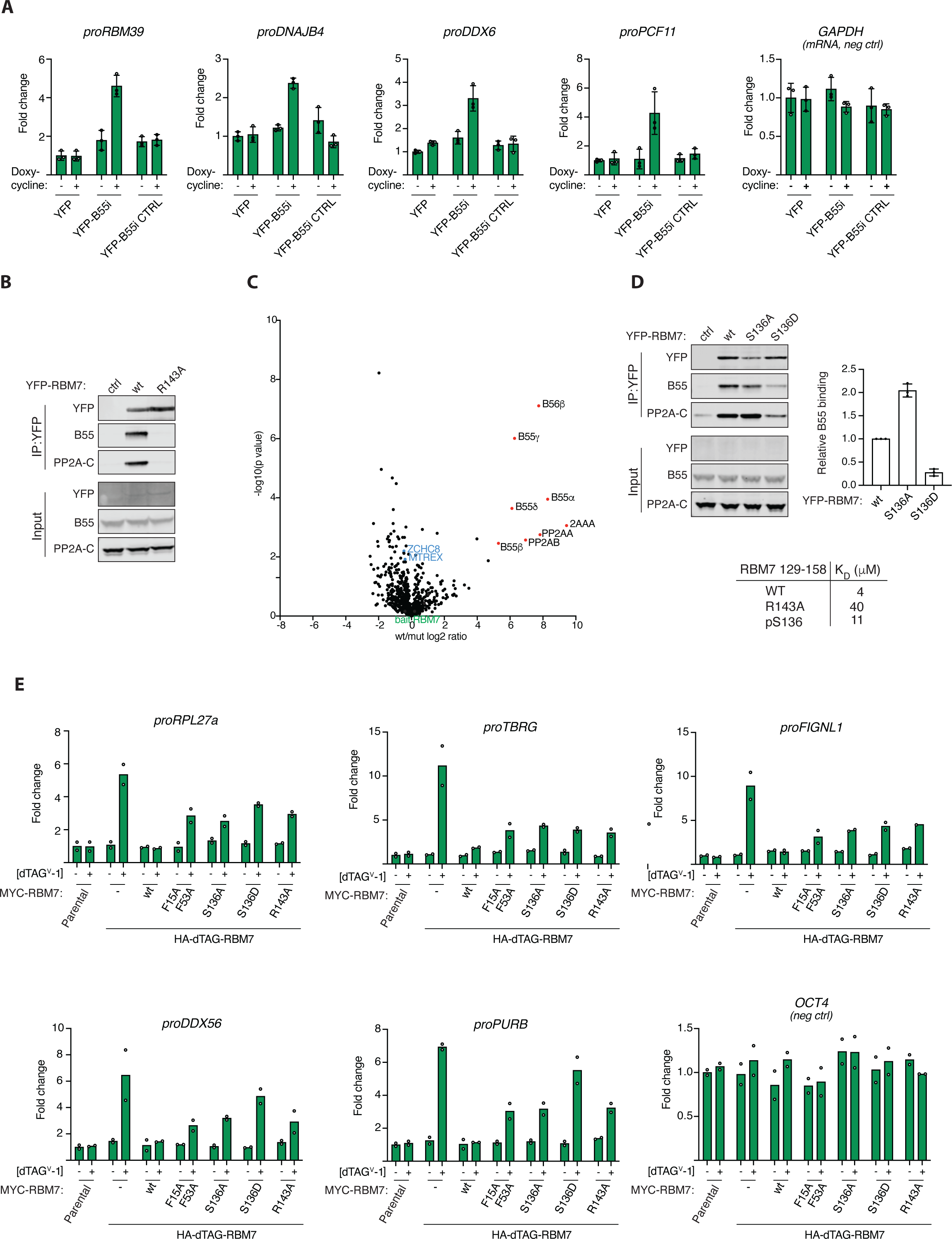
PP2A-B55 regulates NEXT complex function through binding RBM7. **A)** Stable HeLa cell lines expressing the indicated constructs and RNA levels of indicated NEXT substrates measured by RTqPCR. **B**) IP of RBM7 constructs and monitoring binding to B55. **C**) Mass spectrometry comparison of RBM7 WT and R143 with NEXT components in blue. **D**) Binding of RBM7 S136A and S136D to PP2A-B55 and table of affinities measured by SPR measurements of RBM7 peptides. **E**) Endogenous RBM7 was tagged with dTAG allowing rapid removal of RBM7 by adding dTAG and cells were complemented with the endicated RBM7 variants and the indicated RNAs were quantified by RTqPCR.

To explore this in more detail, we turned to our AF modelling of the RBM7-B55 complex. This revealed a critical role of RBM7 R143 in binding to B55, which we confirmed by immunopurifications and by affinity measurements of RBM7 peptides (Fig. 4B, Fig. S16). Comparative mass spectrometry analysis of immunoprecipitations of RBM7 WT and the corresponding R143A variant detected no defect in NEXT complex assembly (ZCCHC8, MTR4/MTREX), but demonstrated the efficient uncoupling of PP2A-B55 binding of RBM7 R143A (Fig. 4C). Moreover, our AF model showed that phosphorylation of S136 in RBM7, a reported functional phosphorylation site (*40*), would negatively regulate PP2A-B55 binding through electrostatic repulsion with acidic residues in B55. We confirmed this in binding assays, suggesting a mechanism for how PP2A-B55 interaction with helical binding elements can be regulated (Fig. 4D and Fig. S16).

To determine if PP2A-B55 binding to RBM7 was required for *in vivo* function of the NEXT complex, we utilized a genetic complementation system in mouse embryonic stem (mES) cells where endogenous RBM7 could be inducibly degraded, using the dTAG system (*42*), which allowed for the functional investigation of stably expressed RBM7 variants (Fig. 4E and S16). As a functional read-out, we monitored levels of a panel of NEXT RNA substrates in RBM7 WT, S136A/D, and R143A complemented cells. As controls we included the RBM7 F15A/F53A variant, with diagnostic point mutations in the RNA-recognition motif (RRM) domain (*43*), and also measured RNA levels of *OCT4* mRNA, which is not a NEXT substrate. While RBM7 WT fully suppressed the increased NEXT substrate levels, arising upon removal of endogenous RBM7, both the RBM7 S136 and R143 mutants failed to do so (Fig. 4E). Thus, regulated binding of PP2A-B55 to RBM7 appears important for NEXT function. We anticipate that phosphorylation sites in RBM7 itself as well as sites in NEXT complex associated factors are dephosphorylated by PP2A-B55 to regulate RNA turnover in response to cellular cues. This is consistent with our observation of changes in RBM7 phosphorylation patterns when uncoupled from PP2A-B55 (Supplemental Fig. S5) and recent phosphoproteomic studies identifying several phosphosites on ZCCHC8 to be targets of PP2A-B55 (*26, 44*).

## Discussion

Here, we uncover how PP2A-B55 binds to cellular binding partners to regulate signaling pathways. A general concept emerging from our work is that PP2A-B55 binds to α-helices in proteins. Despite limited sequence identity, these helices engage many of the same hydrophobic and charged regions on B55. This mechanism of binding is distinct from that of SLiMs, representing a novel mechanism by which PPPs bind substrates. Interestingly, we observe a high degree of flexibility in the composition of binding α-helices. This may allow for them to arise *de novo* through a limited number of mutations similar to how SLiMs can arise *de novo*. Thus, during evolution, PP2A-B55 can readily be integrated into signaling programs to tune their outputs and dynamics. Our work argues that several of the α-helices we identified act as substrate specifiers, a claim that is strongly supported by our ability to turn FOXO3 into a PP2A-B55 substrate by fusing it to the SERTA domain from CDCA4. We anticipate that once bound, PP2A-B55 can dephosphorylate several proteins in the vicinity of its binding site. Interestingly, recent cryo-EM structures of endogenous inhibitors revealed that these also bind this conserved surface of B55 using short α-helices while making additional contacts to the catalytic subunit (*21*). Thus, the function of inhibitors is two-fold in blocking both substrate recruitment and activity.

Inspired by how the endogenous PP2A-B55 inhibitors work we used deep learning protein design to generate a potent and specific B55 inhibitor that can outcompete substrate binding. We show that this inhibitor can be used to dissect PP2A-B55 functions in biological pathways such as cell division and RNA degradation. Our work outlines a general strategy for how *de novo* protein design can generate powerful tools for dissecting biological questions and provide tools for therapeutic proof of principle studies.

Collectively, we provide important insight into how PP2A-B55 achieves binding specificity and a foundation for understanding and precisely engineering PP2A-B55 programs to dissect cellular signaling throughout the eukaryotic domain of life.

## Supporting information

Combined supplemental figures

## Acknowledgements

Work at the Novo Nordisk Foundation Center for Protein Research is supported by NNF14CC0001. Work in the Nilsson lab is supported by grants from the Novo Nordisk Foundation (NNF0082227 and NNF0065098) and the Danish Cancer Society (R269-A15586-B71). We thank the protein production and characterization facility for all their help with the project. J.K.V. was supported by the European Union’s Horizon 2020 UBIMOTIF programme (860517). This work was supported, in whole or in part, by the Israel Science Foundation, founded by the Israel Academy of Science and Humanities (grant number 301/2021 to OS-F). Work in the T.H.J. laboratory was supported by the Independent Research Fund Denmark - Medical Sciences and the Novo Nordisk Foundation (NNF, ExoAdapt Grant 31199). Work in AK lab is supported by NIH-R35GM119455. AAJ was supported by Wellcome Senior Research Fellowship (202811). AAJ and his team are co-funded by the European Union (ERC, CHROMSEG, 101054950). Views and opinions expressed are however those of the author(s) only and do not necessarily reflect those of the European Union or the European Research Council. Neither the European Union nor the granting authority can be held responsible for them.

## Author contribution

TK performed immunoprecipitation experiments and live cell analysis of FOXO3 and B55i. DHG performed immunoprecipitation experiments and prepared samples for mass spectrometry analysis. JG and OSF performed AF modelling and analysis and inhibitor design. WG and THJ planned, performed and analysed NEXT complex experiments. BM and AK performed mass spectrometry analysis. JBH and JN initiated the project and conducted immunoprecipitation experiments. MBW helped with AF modelling. IBP prepared samples for MS. CBC and BLM performed affinity measurements. PSP and JP helped with reconstitution experiments. JN drafted manuscript and coordinated work.

## Methods

### Expression constructs and Immunoprecipitation

Standard cloning techniques were used throughout. Point mutations were introduced by whole plasmid PCR. All constructs were fully sequenced. Synthetic DNA was purchased from GeneArt, Life Technologies and cloned into pcDNA5/FRT/TO (Invitrogen) expression vector containing YFP resulting in the indicated YFP fusion proteins. These constructs were transiently transfected into HeLa cells 24 hours prior to harvesting cells. Cells were lysed in lysis buffer (50 mM Tris-HCl pH 7.5, 50 mM NaCl, 1 mM EDTA, 1 mM DDT and 0.1% NP40). Complexes were immunoprecipitated at 4°C in lysis buffer with GFP-Trap (ChromoTek) and washed in lysis buffer. Precipitated protein complexes were washed three times in lysis buffer, eluted in 2×SDS sample buffer and subjected to Western blotting or mass spectrometry as indicated.

### B55 inhibitor peptide competition for MS analysis

Ten full 15 Cm^3^ dishes with HeLa FRT cells stably expressing Venus-B55 alpha were induced with 10 ng/mL Doxycycline for 24h and collected by trypsinization. Pellets from one dish were lysed in 400 μL lysis buffer (100 mM NaCl, 50 mM tris pH 7.4, 0,1% NP-40, 1mM DTT supplemented with protease and phosphatase inhibitors (Roche). Lysate was sonicated for 10 cycles (30s On, 30s OFF) at 4°C and kept on ice for 20 min and cleared for 45 min 4°C 20000g. Next the lysate was pooled, 200 μL pre-equilibrated GFP-trap beads were added, mixed, and divided into 10 tubes Lo-bind tubes (Eppendorf). Either wt or ctrl/mutant inhibitor was added to a final concentration of 2 μM. Immunoprecipitations were incubated for 1h at 4°C, washed 3 times with 1 mL lysis buffer and eluted in 40 μL 2x LSB buffer.

### Computational methods Screen of binders

Predictions with AlphaFold2-multimer-v2.3 were run between the 256 proteins of the pull-down using the following parameters, using A100 and A30 GPU-s:

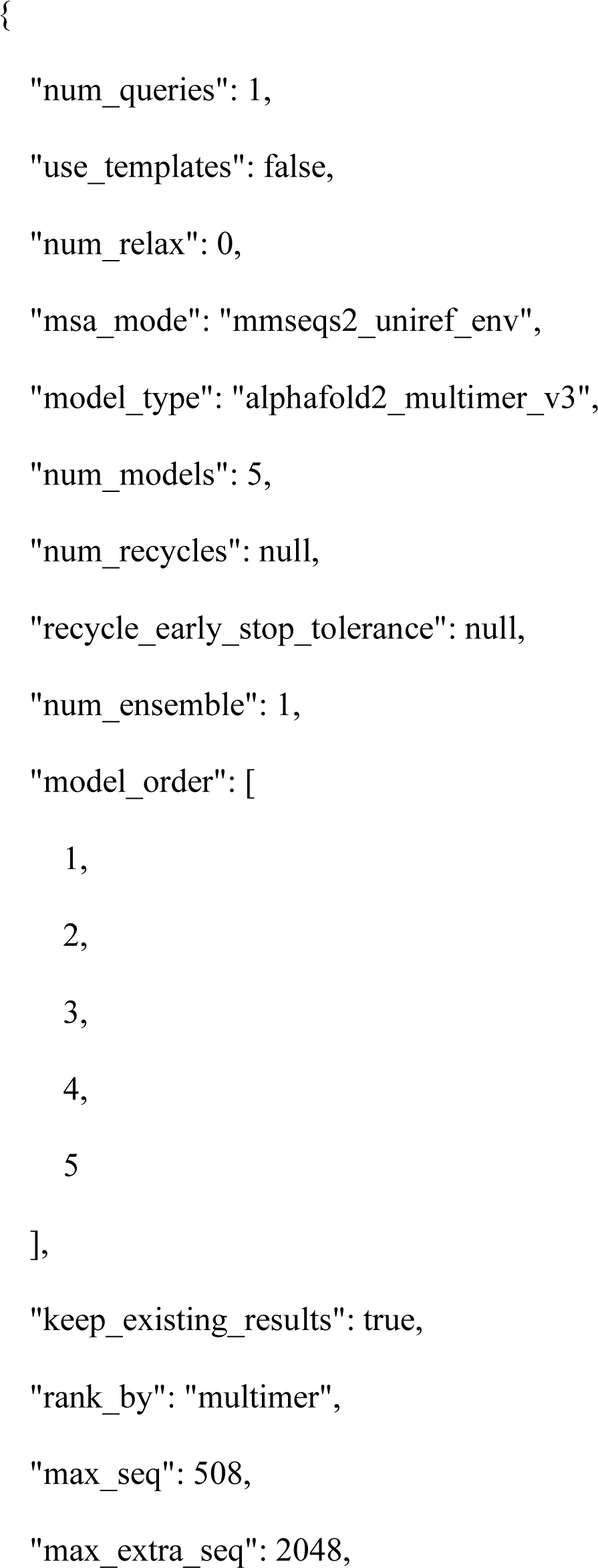
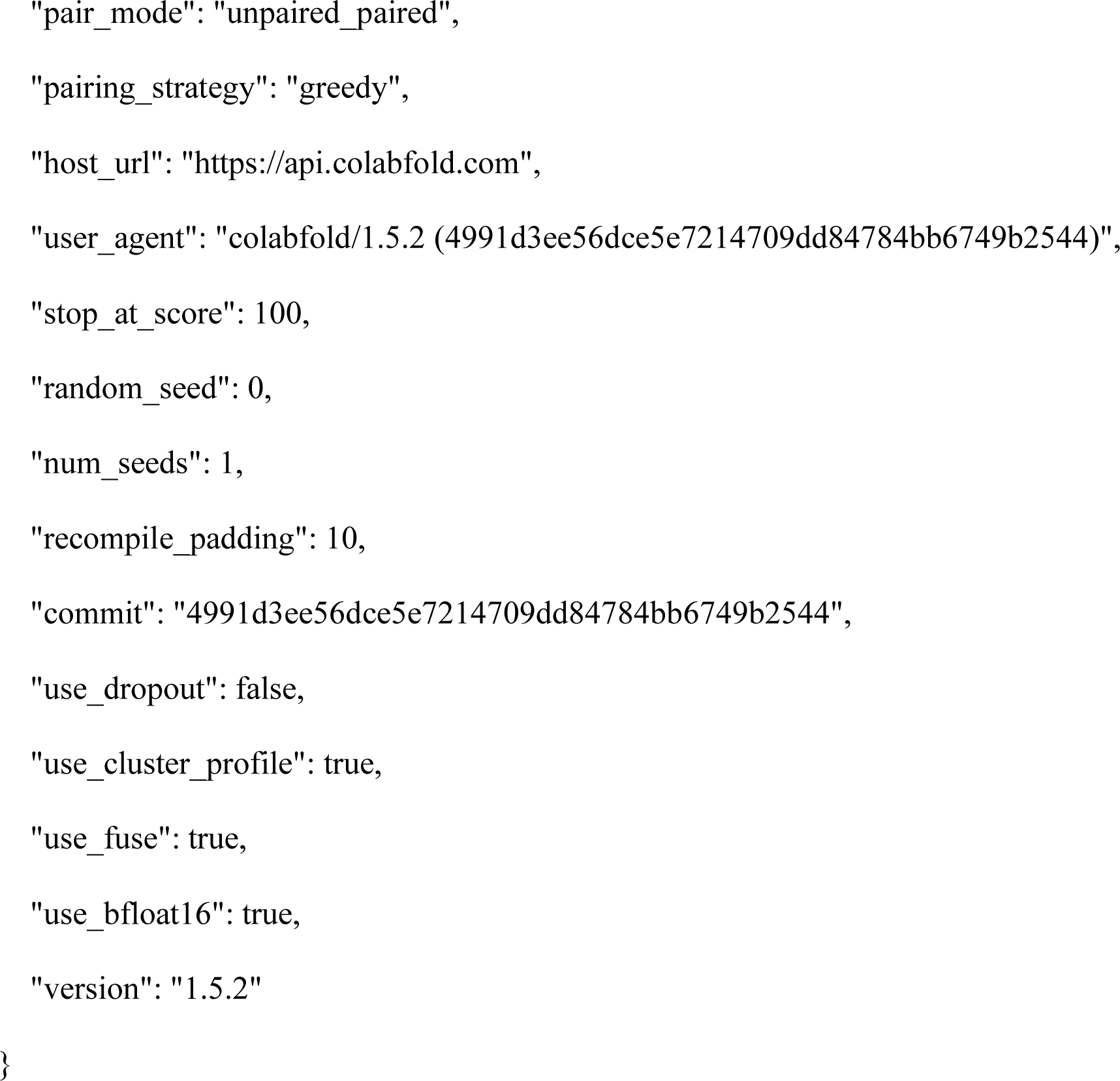

The resulting complexes were filtered by calculating the interface residues between the conserved binding site residues of B55 and the partner. To speed up computation, we first calculated the distance between all Cβ atoms of the two chains and selected those below 8Å, then for these residues, we measured distances between all atoms of the two chains and used the cutoff of 4Å to define interacting residues. Based on our observation that pLDDT confidence of flanking regions around otherwise well predicted regions can drag down the average pLDDT calculated in the next step, we discarded residues with pLDDT below 50 from further calculations. For the remaining interfaces, we calculated the following metrics if the more than seven interface residues remained, otherwise we discarded the model:

- the average pLDDT for the binding partner of B55 over the residues selected previously.
- the iPAE (interface PAE) by selecting the values defined by the peptide (rows) and receptor (columns) interface residues from the PAE matrix and calculating their median.

For models that did not meet these criteria, we set the average pLDDT to 0, and iPAE to 30. We then selected the complexes with iPAE< 10 and average interface pLDDT > 70. Finally, the complexes were manually inspected.

### Design process for de novo binders

The structures of AMOTL2, IER2, CDCA4 and Zds1 were submitted to design with ProteinMPNN (*33*) using the interface at HuggingFace (https://huggingface.co/spaces/simonduerr/ProteinMPNN). 100 sequences were generated for each input template with the default model, temperature 0.1 and backbone noise of 0.3.The chain of B55 was fixed during design and the binder chain was fully designable. The resulting sequences were refolded with AlphaFold2 using local runs with default parameters. Average pLDDT of the binder residues was calculated, as well as the RMSD between the template and the refoled designed binders. Designs with average pLDDT over 90 were manually inspected.

### Visualization of structural models

Structural models were visualized using ChimeraX (*45*).

### Availability of structural models

The AlphaFold2 models are provided as supplemental material.

### Live cell

Live-cell analysis was performed on a Deltavision Elite system using a × 40 oil objective with a numerical aperture of 1.35 (GE Healthcare). The DeltaVision Elite microscope was equipped with a CoolSNAP HQ2 camera (Photometrics). Cells were seeded in eight-well Ibidi dishes (Ibidi) and before filming, the media was changed to Leibovitz’s L-15 (Life Technologies). Appropriate channels were recorded for the times indicated. For transient transfections, DNA constructs were transfected into HeLa cells using jetOPTIMUS reagent (Polyplus) 24 h prior to analysis. The nuclear/cytoplasmic distribution of YFP-FoxO3 was analysed using SoftWoRx (GE Healthcare) software.

### Cell culture

HeLa cells were maintained in DMEM GlutaMAX containing 100 U/ml penicillin, 100 mg/ml streptomycin, and 10% FCS (all from Thermo Fisher Scientific). Stable HeLa cell lines were generated using the T-Rex doxycycline inducible Flp-In system (Invitrogen) and cultivated like HeLa cells with the addition of 5 mg/mL Blasticidin and 100 mg/mL Hygromycin B. *E. coli* DH5alpha were maintained and propagated using standard microbiological procedures. The following drug concentrations were used: Thymidine 2.5 mM and doxycycline 10 ng/ml unless otherwise stated.

### Expression and purification of proteins

B55alpha was cloned into pCPR0197 allowing expression in HEK293 cells as a His-Strep-TEV fusion protein. Following harvesting the cell pellet was resuspended in lysis buffer (100 mM Tris pH=8, 150 mM NaCl, 1mM EDTA, TCEP, protease inhibitors, benzonase) and cells lysed by sonication. Following clarification by centrifugation and filtration the lysate was loaded on a Strep-affinity column and washed with Buffer A (100mM Tris pH=8.0, 150 mM NaCl, 1 mM EDTA, TCEP) and eluted with Buffer A + 2,5 mM desthiobiotin. The peak fractions were pooled and diluted with 100 mM Tris pH=8.0 and loaded on to a MonoQ column. The column was washed with Buffer M (100 mM Tris pH=8.0, 50 mM NaCl, 10% glycerol, TCEP) and eluted with a 50 mM-1M NaCl gradient in Buffer M. Individual fractions were frozen.

His-PME1 was expressed in BL21(DE3) at 18 degrees overnight. Cell pellets were resuspended in Buffer L (50 mM NaP pH=7,5; 300 mM NaCl; 10 mM imidazole; 10% glycerol; 0,5 mM TCEP; protease inhibitors) and lysed by sonication followed by centrifugation to clarify lysate. The lysate was loaded on a His affinity column and washed with buffer W (50 mM NaP pH=7,5; 300 mM NaCl; 30 mM imidazole; 10% glycerol; 0,5 mM TCEP) and eluted with the same buffer but containing 500 mM imidazole. Peak fractions were pooled, concentrated and run on a superdex 200 26/60 equilibrated with Buffer GF(50 mM NaP; 150 mM NaCl; 10% glycerol; 0,5 mM TCEP) and peak fractions pooled.

### SPR

All SPR experiments were performed at 25 °C on a Biacore T200 instrument equipped with CM5 sensor chips (Cytiva, Uppsala, Sweden). SPR running buffers and amine-coupling reagents (N-ethly-N’-(3-dimethlyaminoproply)carbodiimide (EDC), N-hydroxysuccinimide (NHS), and ethanolamine HCl) were purchased from Cytiva. The Twin-Strep-tag® capture kit was from Iba Lifesciences (IBA GmbH, Germany). Peptides were purchased from Peptide 2.0 Inc (Chantilly, USA). 1x PBS-P (11.9 mM NaH_2_PO4-Na_2_HPO4 pH 7.4, 137 mM NaCl, 2.7 mM KCl, 0.005% (v/v) surfactant P20) was used as running buffer for Strep-Tactin®XT immobilization onto flow cells 1 and 2 of the CM5 SPR sensor chip following the protocol from the manufacturer. Subsequently, Twin-Strep tagged B55α at 20 nM was captured on the flow cell 2 (Fc2) channel of the Strep-Tactin®XT coated chip. The flow cell 1 (Fc1) remained unmodified and was used as reference for subtraction of systematic instrumental drift.

Approximately 550 RU of B55α were obtained with a flow rate of 5 μL/min and a contact time of 100 s. The buffer used for B55α capture and the following SPR binding experiments was 20 mM Tris pH 8.0, 250 mM NaCl, 0.5 mM TCEP, 0.05% Tween-20. Peptide and full-length proteins (PPME1, Arpp19 *S62E*) threefold serial dilutions were prepared in the SPR running buffer from the primary stocks and a series of concentrations (2 or 3) run in duplicates. Peptides and full-length proteins (PPME1, Arpp19 *S62E*) were injected sequentially over the two flow cells at a flow rate of 30 μl/min or 60 μl/min (Cdca4-1) for 80 s. The dissociation rate of the complexes was monitored for 480 s or 600 s (Cdca4-1/B55α). All binding experiments were run at least in duplicates to confirm the reproducibility of the assay. Data processing and fitting was done using the BiaEvaluation software (v. 3.2.1, Cytiva, Uppsala, Sweden). The raw sensorgrams were double referenced (referring to the subtraction of the data over the reference surface and the average of the buffer injections from the binding responses). The equilibrium dissociation contants, K_D_, were determined by plotting the equilibrium responses levels (Req) against the analyte concentrations and fitting to a steady-state model. For the Cdca4-1/B55α complex, the association and dissociation phases over all replicates were globally fit using a 1:1 interaction model yielding single values for the *k*_a_ and the *k*_d_. The equilibrium dissociation constant, *K*_D_, is the rate of the *k*_d_ over the *k*_a_.

### Fluorescent polarization assay

The binding affinity between the fluorescent peptide probe (FITC-PEG2-LLELAKKKLKELEEEPDPDLRKKTLVRN-Nle-IKKLEW) and B55α was determined as the K_d_ value by saturation binding experiments, where increasing concentrations of B55α (1.8-500 nM) were added to a fixed concentration of peptide probe (3 nM). The assay was performed in a 1 × HBS-T buffer (10 mM HEPES, 150 mM NaCl, 0.005% Tween20, pH = 7.4) using black flat-bottom 384-well plates (Corning Life Sciences, NY), and a volume of 30 μL/well. The assay plate was spun down to ascertain proper mixing and removal of potential air bubbles before measuring the FP levels on a Safire2 plate reader (Tecan, Mannedorf, Switzerland). The g-factor was adjusted at each experiment so that a series of three blank wells containing probe but no B55α defined the baseline FP value. The probe was measured at an excitation/emission value of 470:535 nm. The FP values were fitted to the one-site specific binding equation: Y = B_max_ × X/(K_d_ + X), with B_max_ being the maximal FP value, X the PP2A B55α concentration, and Y the variable FP values. The K_d_ values were derived from the resulting binding saturation curve as being equal to the B55α concentration, where the curve is half-saturated.

### Mass spectrometry

Pull-downs were analyzed on a Q-Exactive Plus quadrupole or Fusion Orbitrap Lumos mass spectrometer (ThermoScientific) equipped with Easy-nLC 1000 or 12000 (ThermoScientific) and nanospray source (ThermoScientific). Peptides were resuspended in 5% methanol / 1% formic acid and loaded onto a trap column [1 cm length, 100 μm inner diameter, ReproSil, C18 AQ 5 μm 120 Å pore (Dr. Maisch, Ammerbuch, Germany)] vented to waste via a micro-tee and eluted across a fritless analytical resolving column (35 cm length, 100 μm inner diameter, ReproSil, C18 AQ 3 μm 120 Å pore) pulled in-house (Sutter P-2000, Sutter Instruments, San Francisco, CA) with a 45-min gradient of 5–30% LC-MS buffer B (LC-MS buffer A: 0.0625% formic acid, 3% ACN; LC-MS buffer B: 0.0625% formic acid, 95% ACN - Q-Exactive Plus quadrupole or: 0.0625% formic acid, 80% ACN Fusion Orbitrap Lumos mass).

Raw data were searched using COMET (*46*)(release version 2014.01) in high-resolution mode against a target-decoy (reversed) (*47*) version of the human proteome sequence database (UniProt; downloaded 2/2020, 40704 entries of forward and reverse protein sequences) with a precursor mass tolerance of +/- 1 Da and a fragment ion mass tolerance of 0.02 Da, and requiring fully tryptic peptides (K, R; not preceding P) with up to three mis-cleavages or no enzyme for Proteinase K digests. Static modifications included carbamidomethylcysteine and variable modifications included: oxidized methionine and STY phosphorylation. Searches were filtered using orthogonal measures including mass measurement accuracy (+/- 3 ppm), Xcorr for charges from +2 through +4, and dCn targeting a <1% FDR at the peptide level. Quantification of LC-MS/MS spectra was performed using MassChroQ (*48*) and the iBAQ method (*49*). All data was analyzed using R 4.3.0. Missing values were imputed from a normal distribution. To be included in further analysis, proteins had to be identified with more than 1 total peptide and quantified in 2 or more replicates of each sample. In each dataset, the abundance of the protein of interest was normalized to be equal across all samples. Statistical analysis were done by two-tailed Student’s t-test.

### mES cell culture and cell line generation

Mouse embryonic stem (mES) cell lines were descendants of the parental E14TG2a cell line (male genotype, XY). mES cells were cultured on 0.2% gelatin coated plates in 2i/LIF containing medium (1:1 mix of DMEM/F12 (Givco) and Neurobasal (Gibco) supplemented with 1x Pen-Strep (Gibco), 2 µM Glutamax, 50 µM beta-mercaptoethanol (Gibco), 0.1 mM Non-Essential Amino Acids (Gibco), 1 mM sodium pyruvate (Gibco), 0.5x N2 Supplement (Gibco), 0.5x B27 Supplement (Gibco), 3 µM GSK3-inhibitor (CHIR99021), 1 µM MEK-inhibitor (PD0325901) and Leukemia Inhibitory Factor (LIF, produced in house) at 37°C, 5% CO_2_. Cells were passaged every 48-72 hours by aspirating medium, dissociating cells with 0.05% Trypsin-EDTA (Gibco) briefly at 37°C before neutralizing with an equal volume of 1x Trypsin Inhibitor (Sigma) and gentle disruption by pipetting. Cells were pelleted by centrifugation to remove Trypsin before resuspending in 2i/LIF medium and plating ∼ 8×10^4^ cells/ml. CRISPR/Cas9 mediated genomic knock-ins (KI) of N-terminal 2xHA-FKBP-V(dTAG) tags were carried out using homology dependent repair (HDR) donor vectors. HDR vectors were cloned to contain gene specific 5’ and 3’ homology arms (∼500 bp) amplified from WT mES cell genomic DNA and cloned into pGNT vectors along with either HYG-P2A-2xHA-FKBP-V or PUR-P2A-2xHA-FKBP-V tagging cassettes. sgRNAs targeting the TSS of genomic loci were cloned into pSLCas(BB)-2A-GFP vectors (pX458 Addgene plasmid ID: 48138) as previously described (*50*). Cells were transfected using Lipofectamine 3000 (Thermo) with 2 donor plasmids harbouring distinct selection markers (HYG/PUR) along with a sgRNA/Cas9 vector in a 1:1:1 ratio. Cells were maintained under double HYG/PUR selection to increase the likelihood of homozygous knock-in clones. Single cell clones were expanded and screened by western blotting analysis before confirming genomic integrations by sanger sequencing of the target locus.

### cDNA cloning and exogenous expression of RBM7

Mouse RBM7 cDNA constructs were cloned using a full-length mouse pUC[mRBM7] cDNA plasmid (Sino Biological, MG5399-U) as a template. RBM7 constructs were amplified by Phusion DNA polymerase (NEB) using standard conditions and mutations were introduced in primer sequences. Fragments were cloned into piggyBAC (pB) vectors harbouring a N-terminal MYC tag and blasticidin (BSD) resistance selection marker using NEBuilder HiFi DNA assembly. HA-dTAG-RBM7 cells were transfected with pB[MYC-RBM7^x^] BSD plasmids along with a pB transposase expressing vector (pBASE) using Viafect transfection reagent (Promega). Cell pools were selected using BSD for 7-10 days or until negative control cells no longer survived. Expression of constructs were validated by western blotting analysis using MYC antibodies.

### Western blotting

Whole cell protein lysates were prepared using RSB100 lysis buffer (10 mM Tris-HCl pH 7.5, 100 mM NaCl, 2.5 mM MgCl_2_, 0.5% NP-40, 0.5% Triton X-100) freshly supplemented with protease inhibitors (Roche). Samples were denatured by the addition of NuPAGE Loading buffer (Invitrogen) and NuPAGE Sample Reducing Agent (Invitrogen) before boiling at 95°C for 5 minutes. SDS-PAGE was carried out on NuPAGE 4-12% Bis-Tris gels (Invitrogen). Western blotting analysis was carried out using standard protocols with the antibodies listed in Table_1 and HRP-conjugated secondary antibodies (Agilent). Bands were visualised bu SuperSignal West Fempto ECL substrate (Thermo) and captured using an Amersham ImageQuant 800 imaging system (GE Healthcare). Images were processed using ImageJ(v.1.53) (*51*).

**Table 1:**
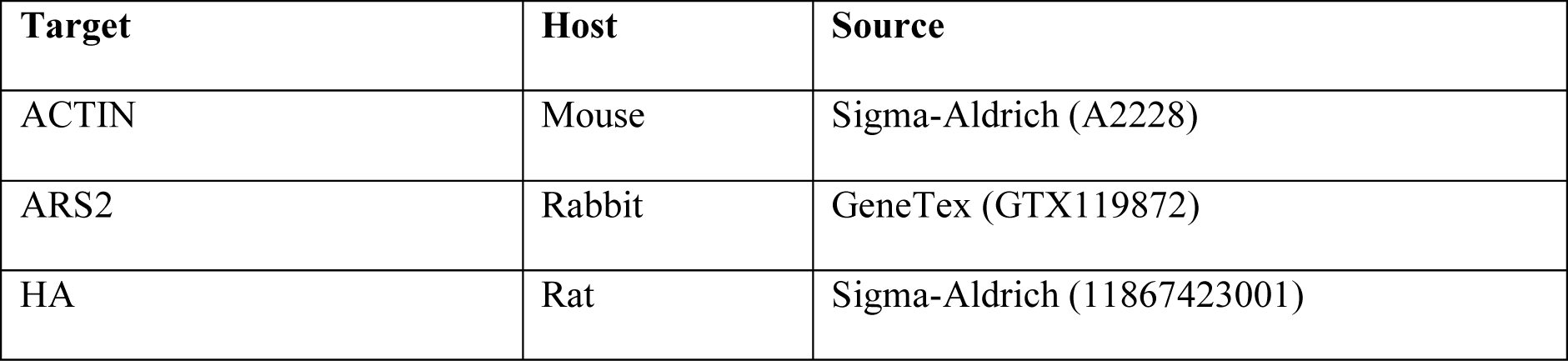
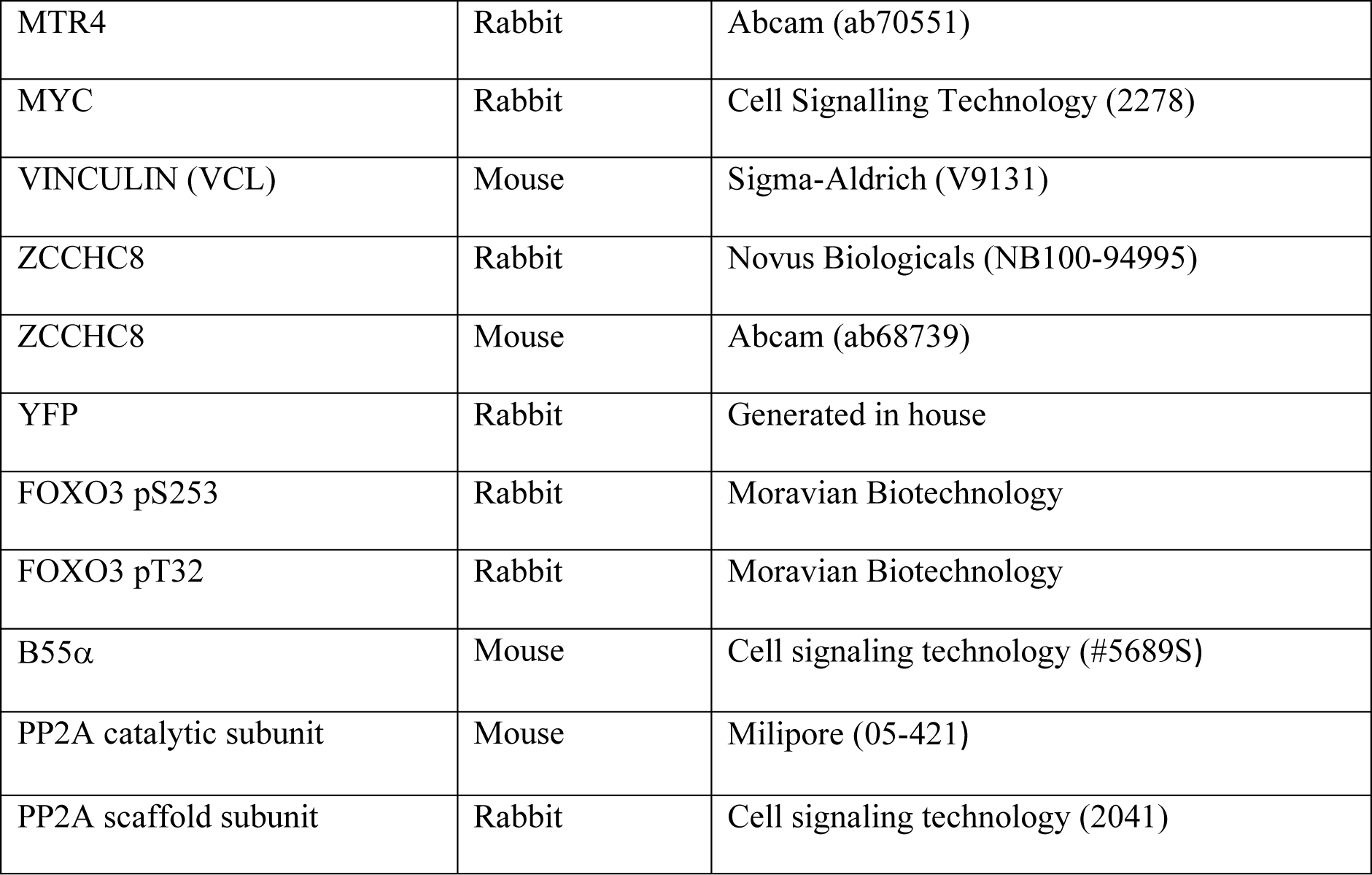
Antibodies used in this study.

### RNA isolation and RTqPCR analysis

Total RNA was isolated using TRIzol (Invitrogen) according to the manufacturer’s instructions. RNA extracts were treated with TURBO DNase (Invitrogen) according to the manufacturer’s instructions, followed by cDNA synthesis from 2 µg total RNA using SuperScript III reverse transcriptase (Invitrogen) and a mixture of 80 pmol random primers (Invitrogen) and 20 pmol oligo d(T)20VN (Merck). qPCR was performed using Platinum SYBR Green (Invitrogen) and an AriaMX Real-Time PCR machine (Agilent). Primers used for RTqPCR are listed in Table_2.

**Table 2:**
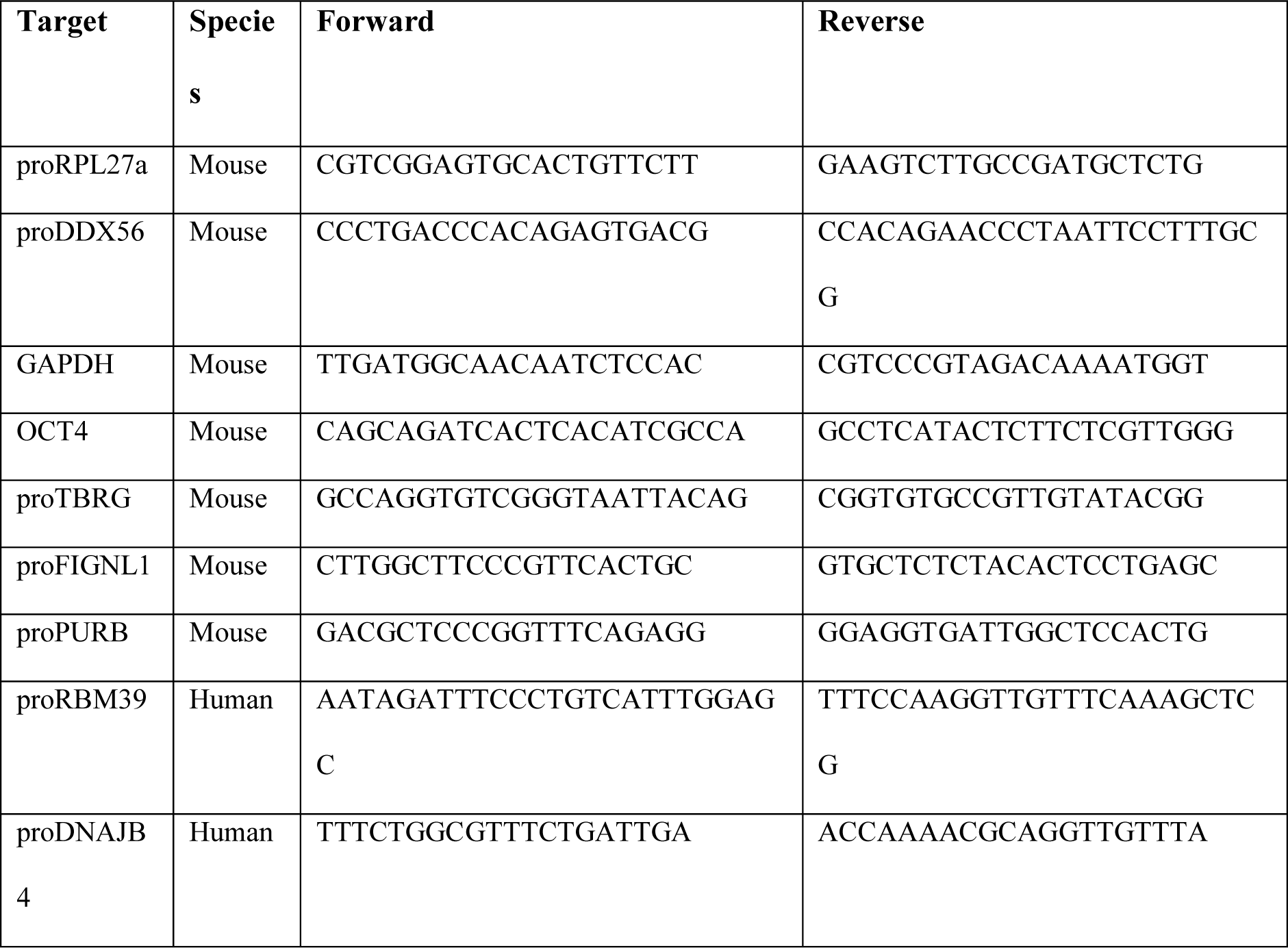
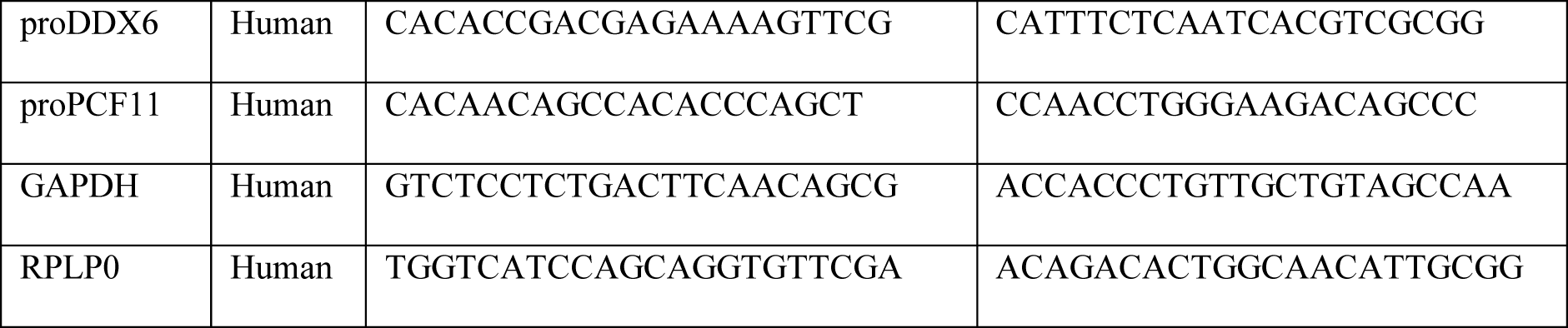
RTqPCR primers used in this study.

