## Supplementary material for "Substrate recognition principles for the PP2A-B55 protein phosphatase": Combined supplemental figures

**Overview of supplemental materials**

Table S1: AF model scores
Table S2: Mass spectrometry data
Supplemental dataset 1:alphafold models pdb files.

Figure S1: AF score plots and comparison to cryo-EM models.
Figure S2: Validation of novel binders and AF models of distinct binding mechanisms.
Figure S3: CDCA4 data.
Figure S4: IER2 data.
Figure S5: RBM7 data.
Figure S6: PME1 data.
Figure S7: EYA3 data.
Figure S8: AMOTL2 data.
Figure S9: Zds1 data.
Figure S10: E4ORF4 data.
Figure S11: Conservation of SERTA domains.
Figure S12: Volcano plots of MS data.
Figure S13: B55 mutant IPs with CDCA4.
Figure S14: Protein MPNN scores in relation to binding.
Figure S15: SPR data for PME1, Arpp19 and RBM7.
Figure S16: Western blots relating to RBM7 biology and RBM7 SPR data.

**A**

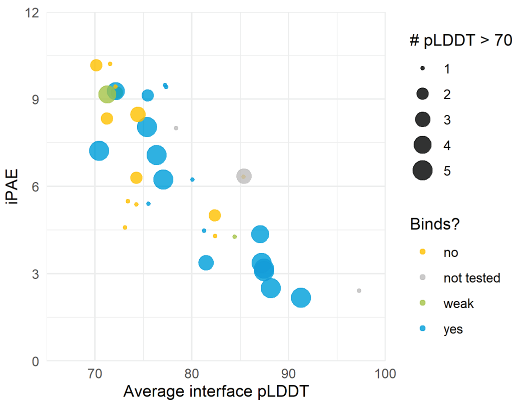


**B**

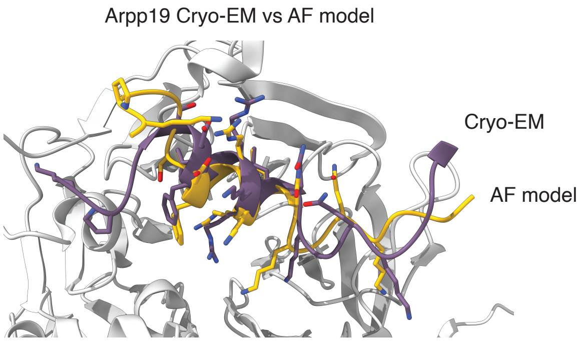


**C**

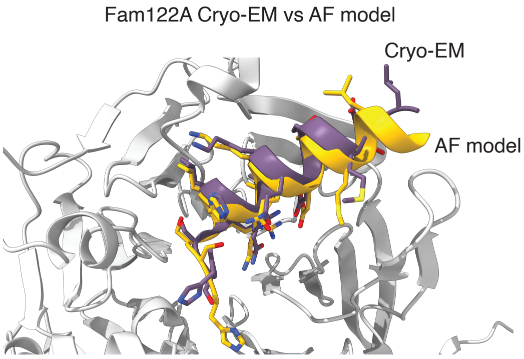


**Supplemental Figure S1.**
**A)** Plot showing iPAE and pLDDT score for the 36 models and whether they were binding to PP2A-B55 in immunoprecipitation experiments. The size of the dot indicates number of models with a pLDDT above 70. **B)** Overlay of the Arpp19 cryo-EM structure (PDB: 8TTB) with the AF model focusing on the helix binding B55. **C)** Overlay of the FAM122A cryo-EM structure (PDB: 8SO0) with the AF model focusing on the helix binding B55.

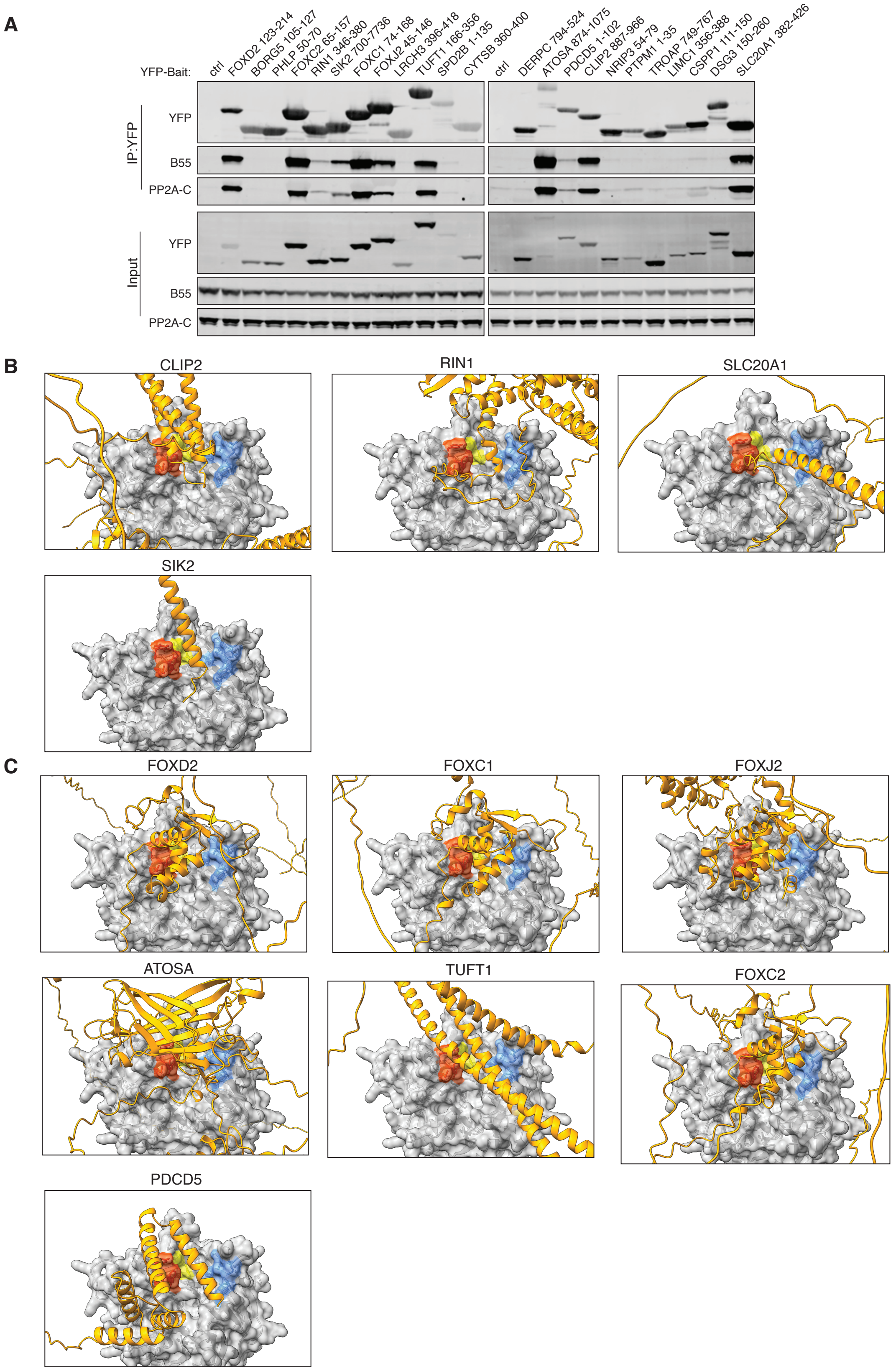


**Supplemental Figure S2**

**A)** The indicated YFP tagged protein fragments were expressed in HeLa cells and following immunoprecipitation they were probed for binding to PP2A-B55 by western blot. **B)** AF models of the indicated proteins binding to PP2A-B55 in a common manner. **C)** AF models of the indicated proteins binding to PP2A-B55 in a distinct manner.

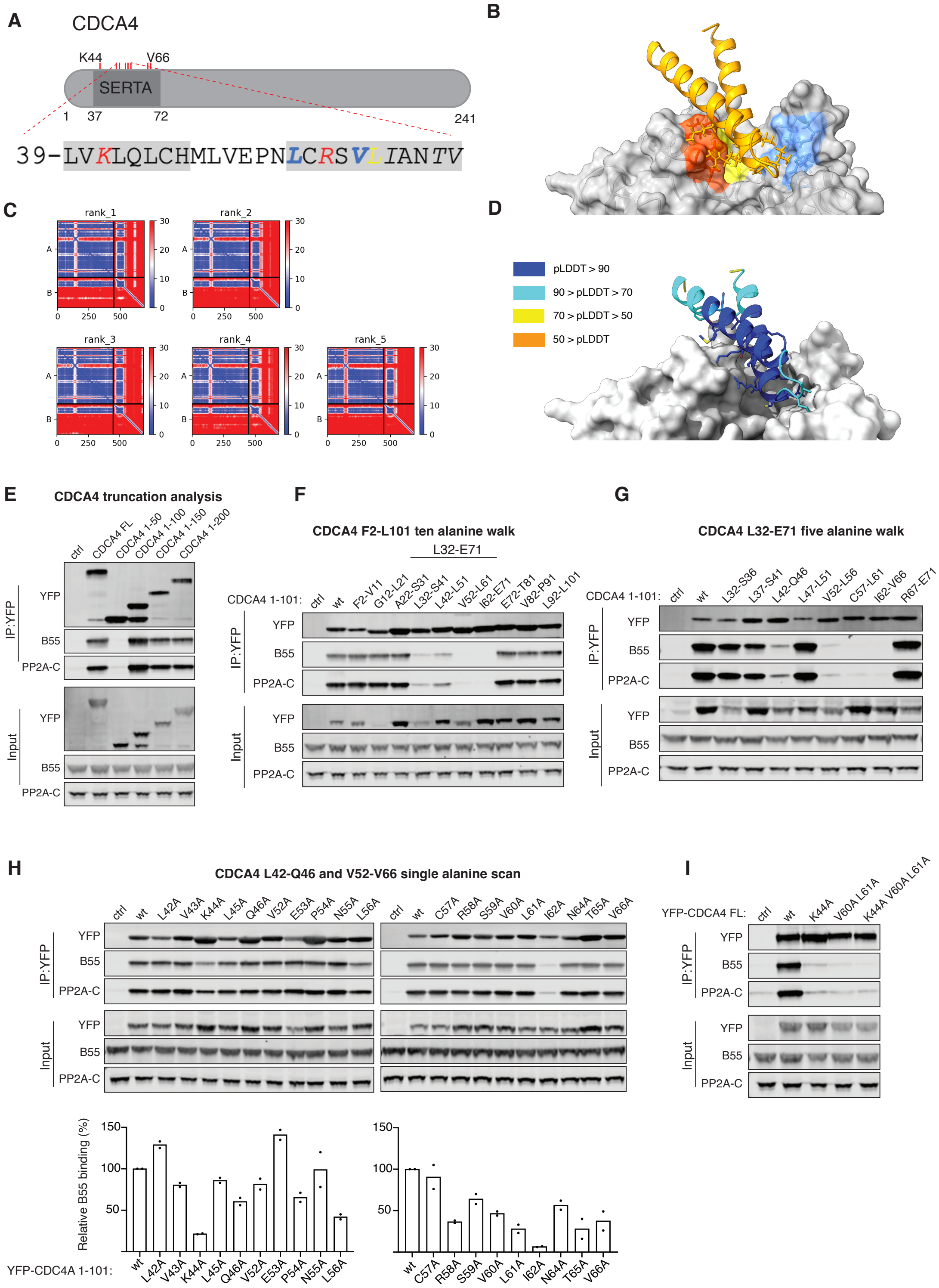


### Supplemental Figure S3

**A)** Schematic of CDCA4 and sequence binding PP2A-B55. **B**) AF model of CDCA4 bound to B55. **C**) Prediction aligned error (PAE) score for the 5 models. **D**) pLDDT score mapped onto model. **E-I**) The indicated YFP tagged protein fragments were expressed in HeLa cells and following immunoprecipitation they were probed for binding to PP2A-B55 by western blot. In H) the quantification of two experiments is shown with binding normalized to wild type protein.

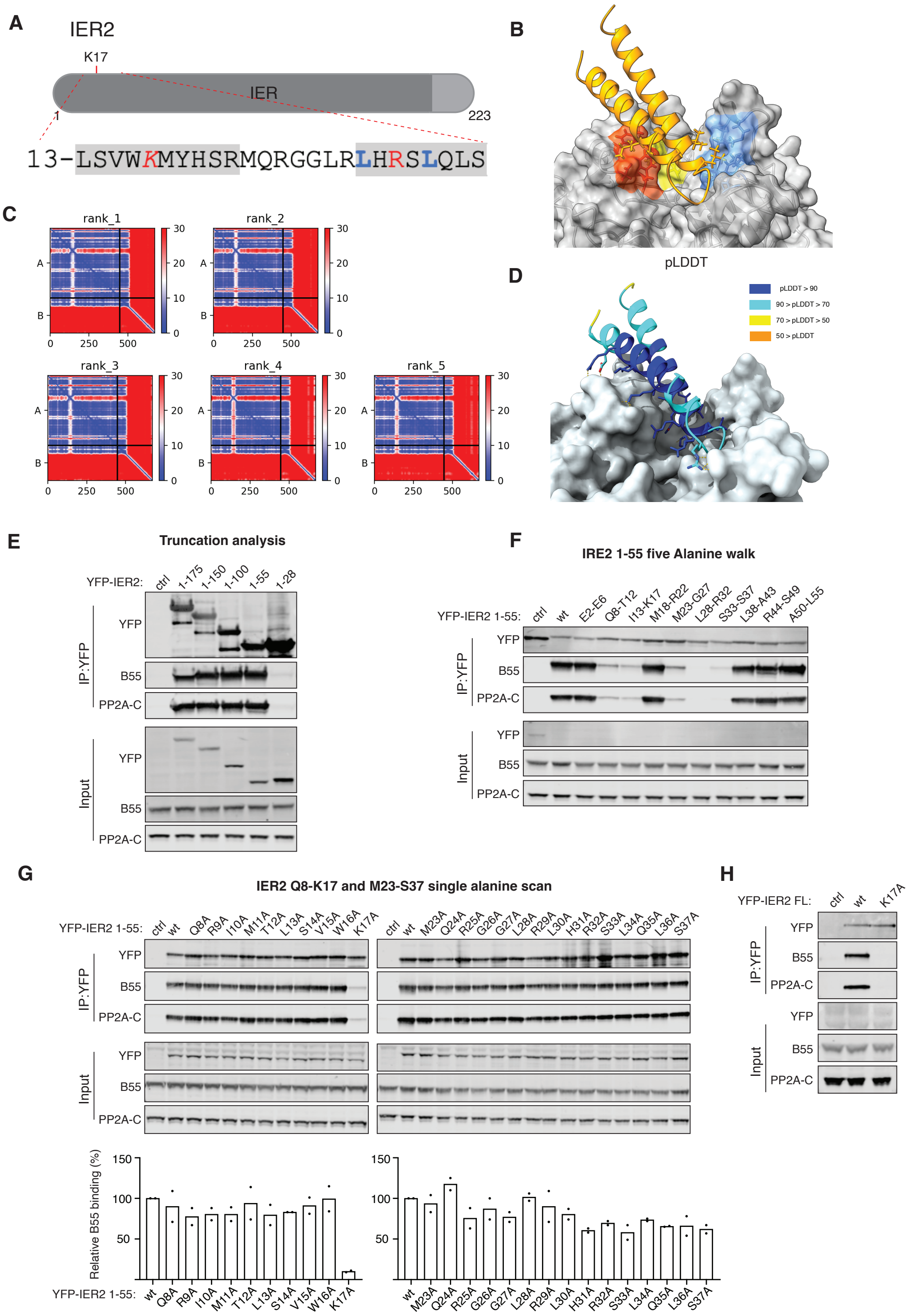


### Supplemental Figure S4

**A)** Schematic of IER2 and sequence binding PP2A-B55. **B**) AF model of IER2 bound to B55. **C**) Prediction aligned error (PAE) score for the 5 models. **D**) pLDDT score mapped onto model. **E-H**) The indicated YFP tagged protein fragments were expressed in HeLa cells and following immunoprecipitation they were probed for binding to PP2A-B55 by western blot.

In G) the quantification of two experiments is shown with binding normalized to wild type protein.

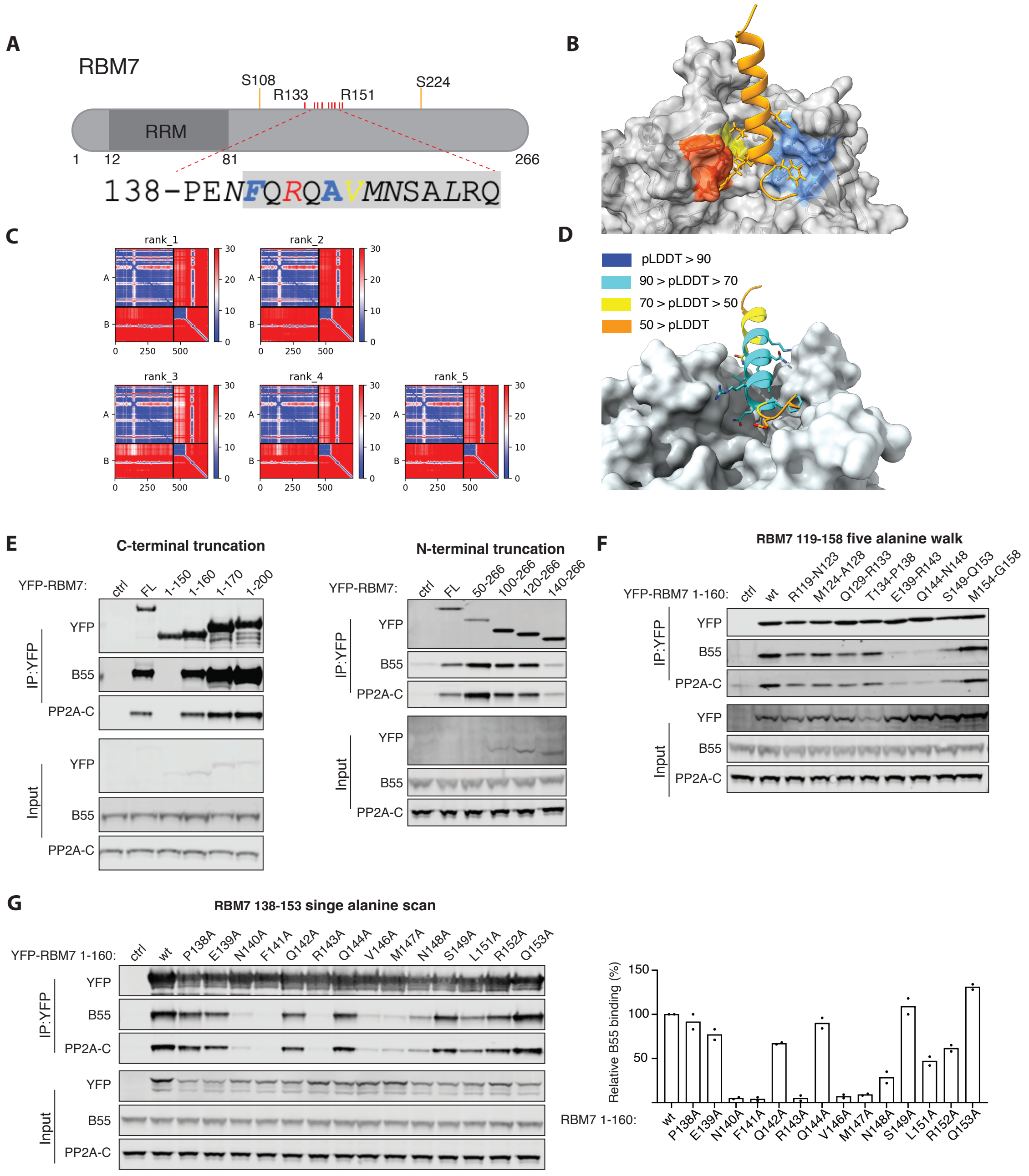


**Supplemental Figure S5**

**A)** Schematic of RBM7 and sequence binding PP2A-B55. Phosphorylation sites with higher occupancy in RBM7 R143A indicated. **B**) AF model of RBM7 bound to B55. **C**) Prediction aligned error (PAE) score for the 5 models. **D**) pLDDT score mapped onto model. **E-H**) The indicated YFP tagged protein fragments were expressed in HeLa cells and following immunoprecipitation they were probed for binding to PP2A-B55 by western blot. In H) the quantification of two experiments is shown with binding normalized to wild type protein.

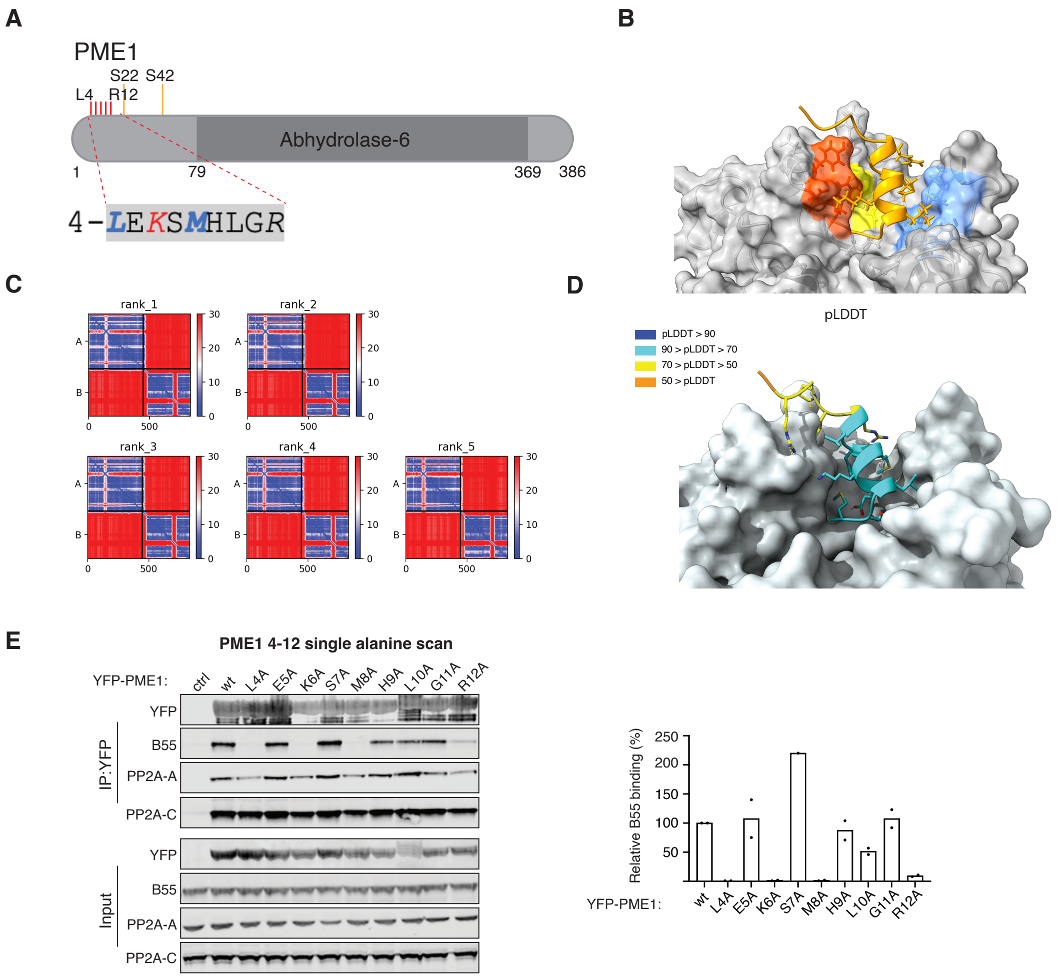


**Supplemental Figure S6**

**A)** Schematic of PME1 and sequence binding PP2A-B55. Phosphorylation sites with higher occupancy in PME1 L4A indicated. **B**) AF model of PME1 binding helix bound to B55. **C**) Prediction aligned error (PAE) score for the 5 models. **D**) pLDDT score mapped onto model. **E**) The indicated YFP tagged proteins were expressed in HeLa cells and following immunoprecipitation they were probed for binding to PP2A-B55 by western blot. The quantification of two experiments is shown with binding normalized to wild type protein.

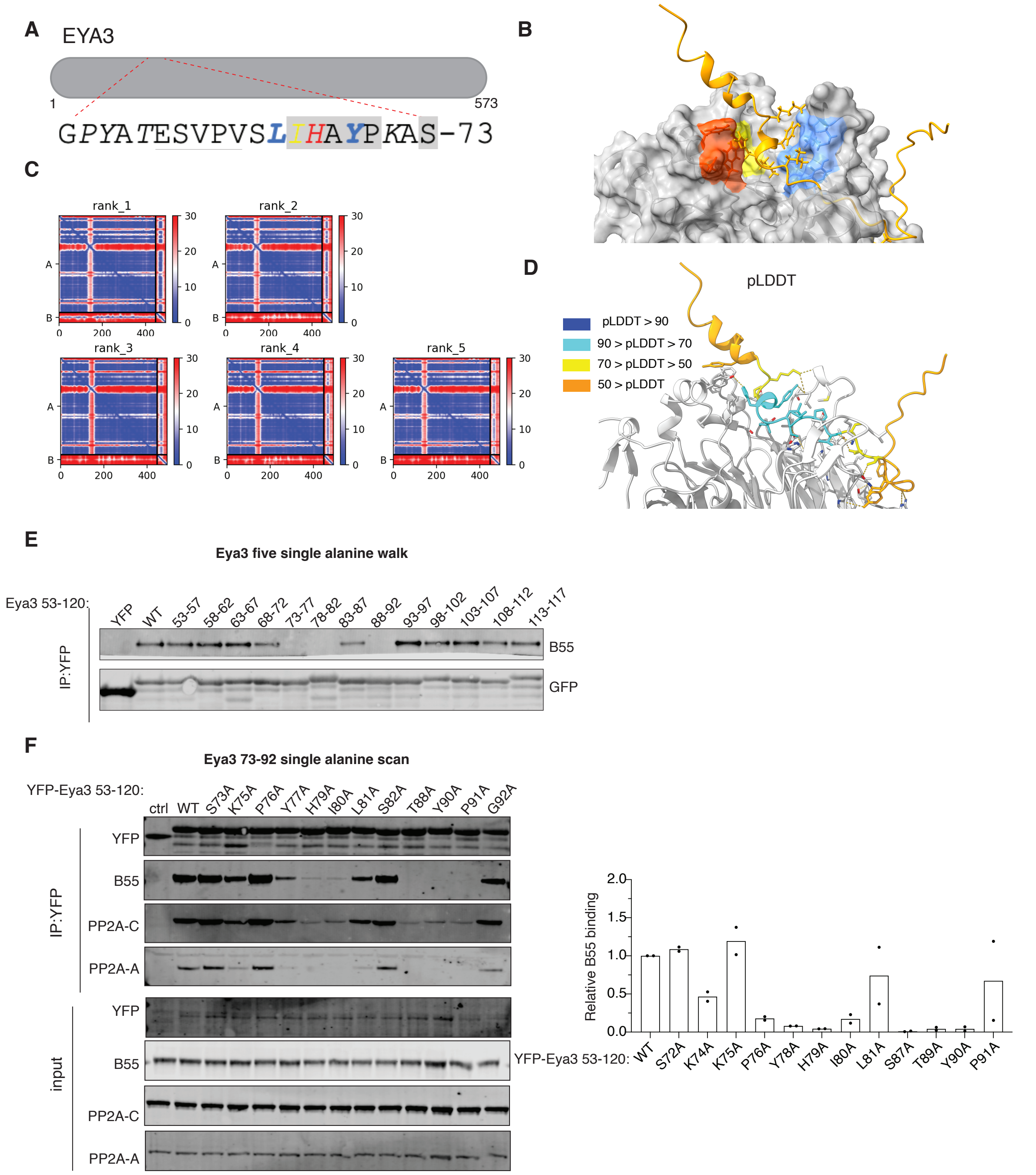


**Supplemental Figure S7**

**A)** Schematic of EYA3 and sequence binding PP2A-B55. **B**) AF model of EYA binding helix bound to B55. **C**) Prediction aligned error (PAE) score for the 5 models. **D**) pLDDT score mapped onto model. **E-F**) The indicated YFP tagged protein fragments were expressed in HeLa cells and following immunoprecipitation they were probed for binding to PP2A-B55 by western blot. In F) the quantification of two experiments is shown with binding normalized to wild type protein.

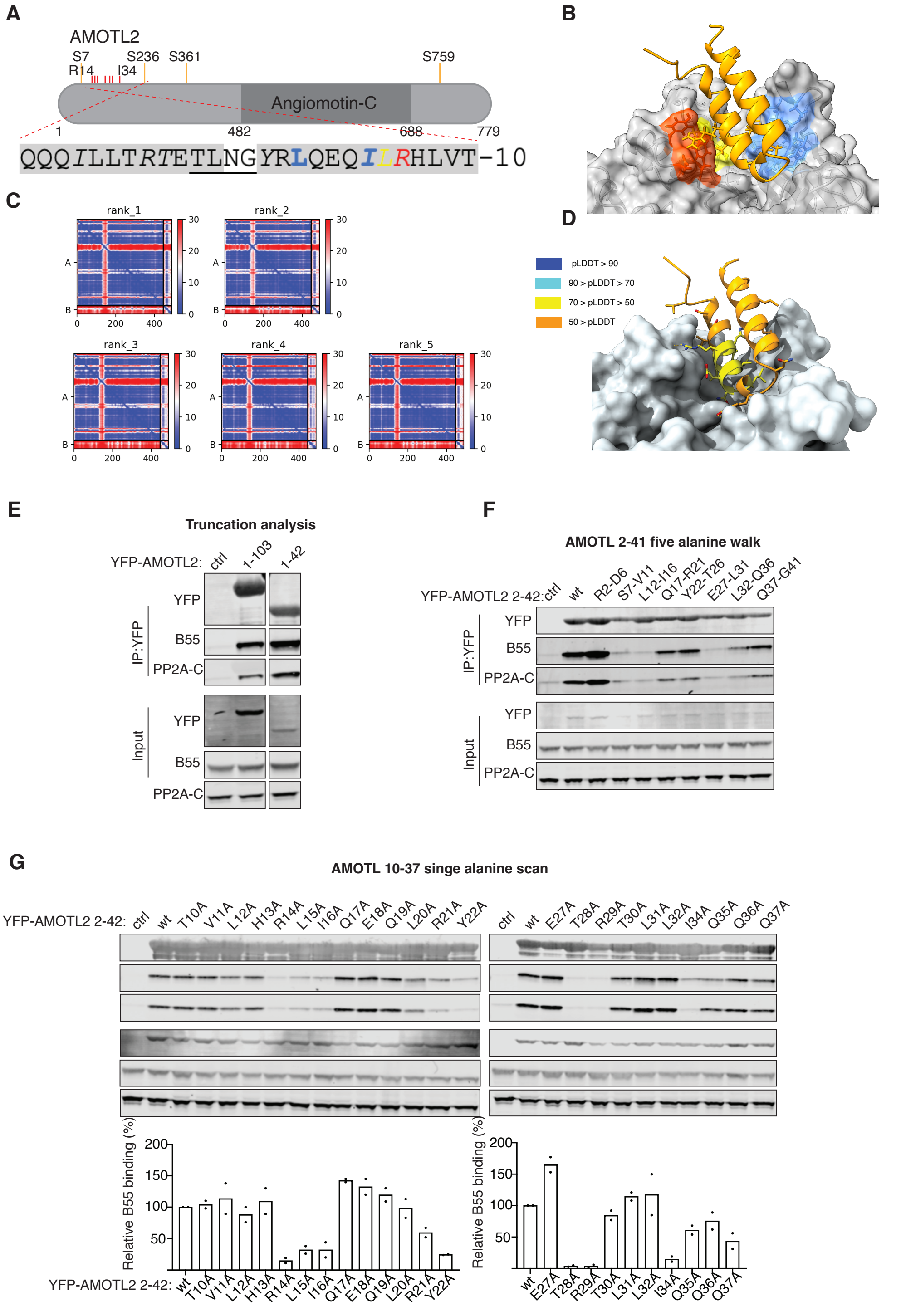


**Supplemental Figure S8**

**A)** Schematic of AMOTL2 and sequence binding PP2A-B55. Phosphorylation sites with higher occupancy in AMOTL2 T28A/R29A indicated **B**) AF model of AMOTL2 bound to B55. **C**) Prediction aligned error (PAE) score for the 5 models. **D**) pLDDT score mapped onto model. **E-I**) The indicated YFP tagged protein fragments were expressed in HeLa cells and following immunoprecipitation they were probed for binding to PP2A-B55 by western blot. In H) the quantification of two experiments is shown with binding normalized to wild type protein.

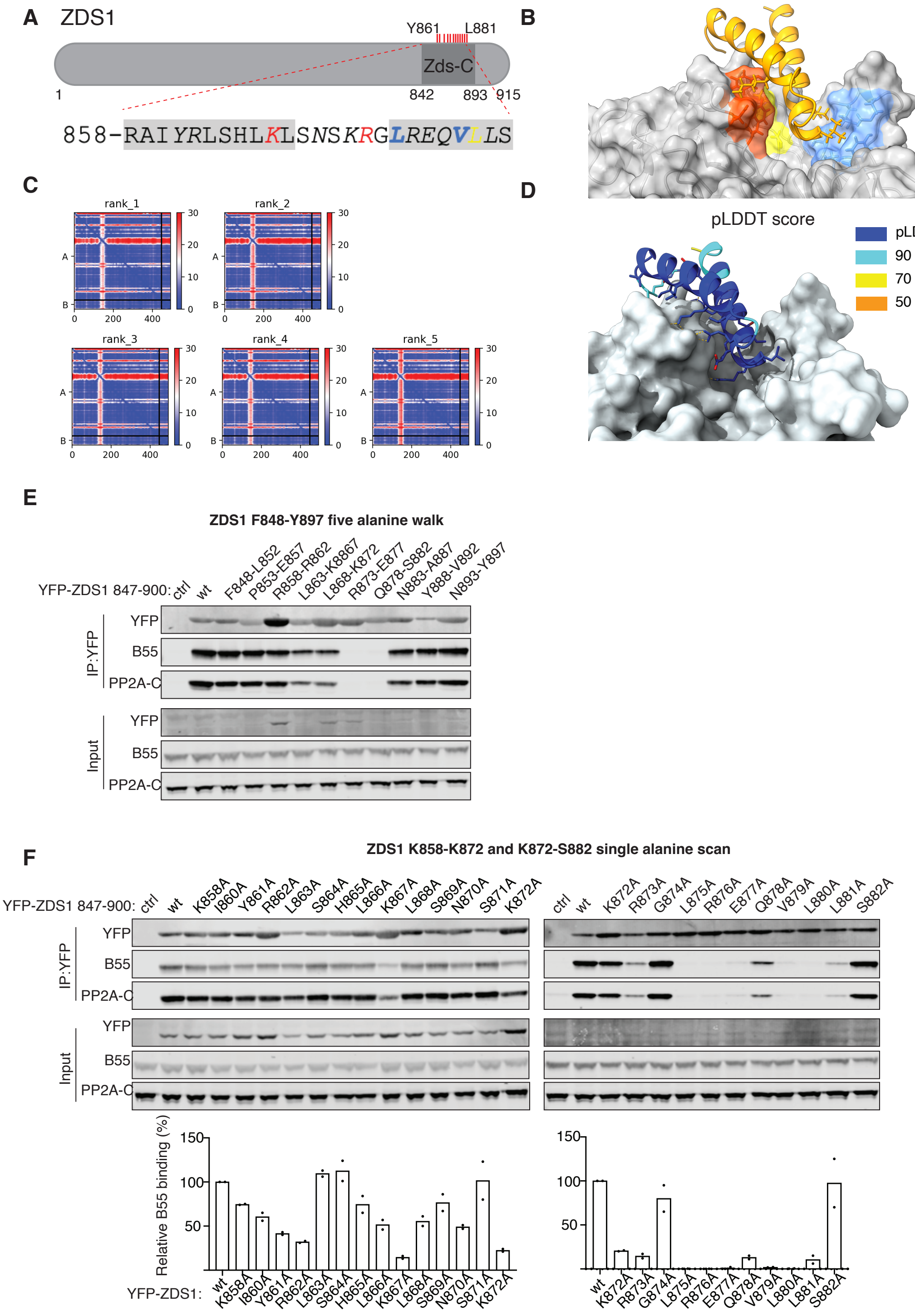


**Supplemental Figure S9**

**A)** Schematic of Zds1 and sequence binding PP2A-B55. **B**) AF model of Zds1 binding helixes bound to B55. **C**) Prediction aligned error (PAE) score for the 5 models. **D**) pLDDT score mapped onto model. **E-F**) The indicated YFP tagged protein fragments were expressed in HeLa cells and following immunoprecipitation they were probed for binding to PP2A-B55 by western blot. In F) the quantification of two experiments is shown with binding normalized to wild type protein.

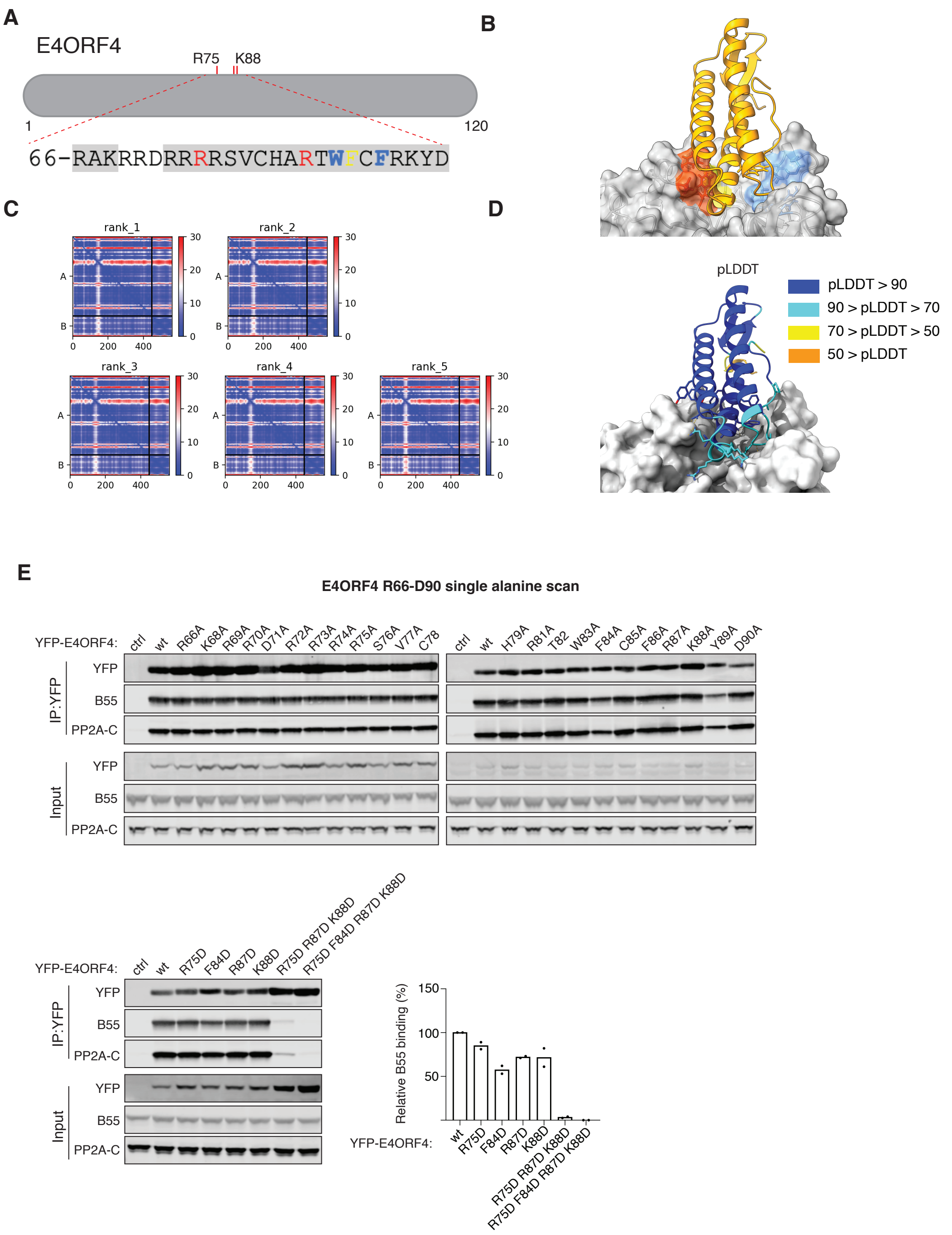


**Supplemental Figure S10**

**A)** Schematic of E4ORF4 and sequence binding PP2A-B55. **B**) AF model of E4ORF4 bound to B55. **C**) Prediction aligned error (PAE) score for the 5 models. **D**) pLDDT score mapped onto model. **E)** The indicated YFP tagged protein fragments were expressed in HeLa cells and following immunoprecipitation they were probed for binding to PP2A-B55 by western blot. **E)** Bottom panel the quantification of two experiments is shown with binding normalized to wild type protein.

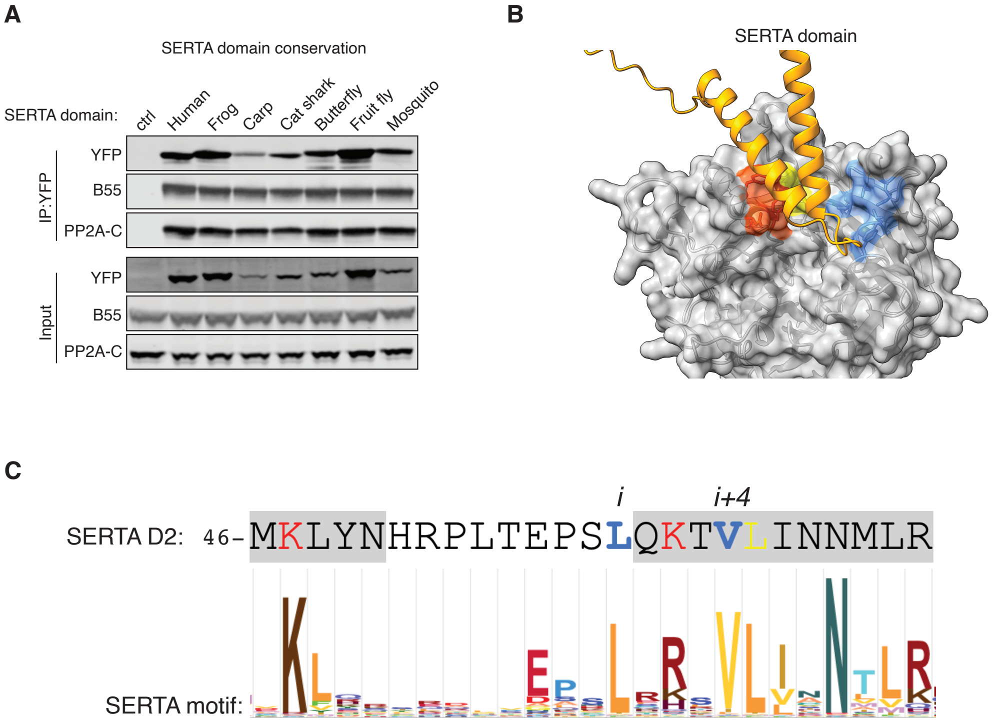


**Supplemental Figure S11**

**A)** YFP tagged SERTA domains from different species were expressed in HeLa cells and binding to PP2A-B55 determined by western-blot. **B**) AF model of the CDCA4 SERTA domain bound to B55. **C**) Signature of all SERTA domains in the pfam database showing conservation of key residues (data from pfam 06031)

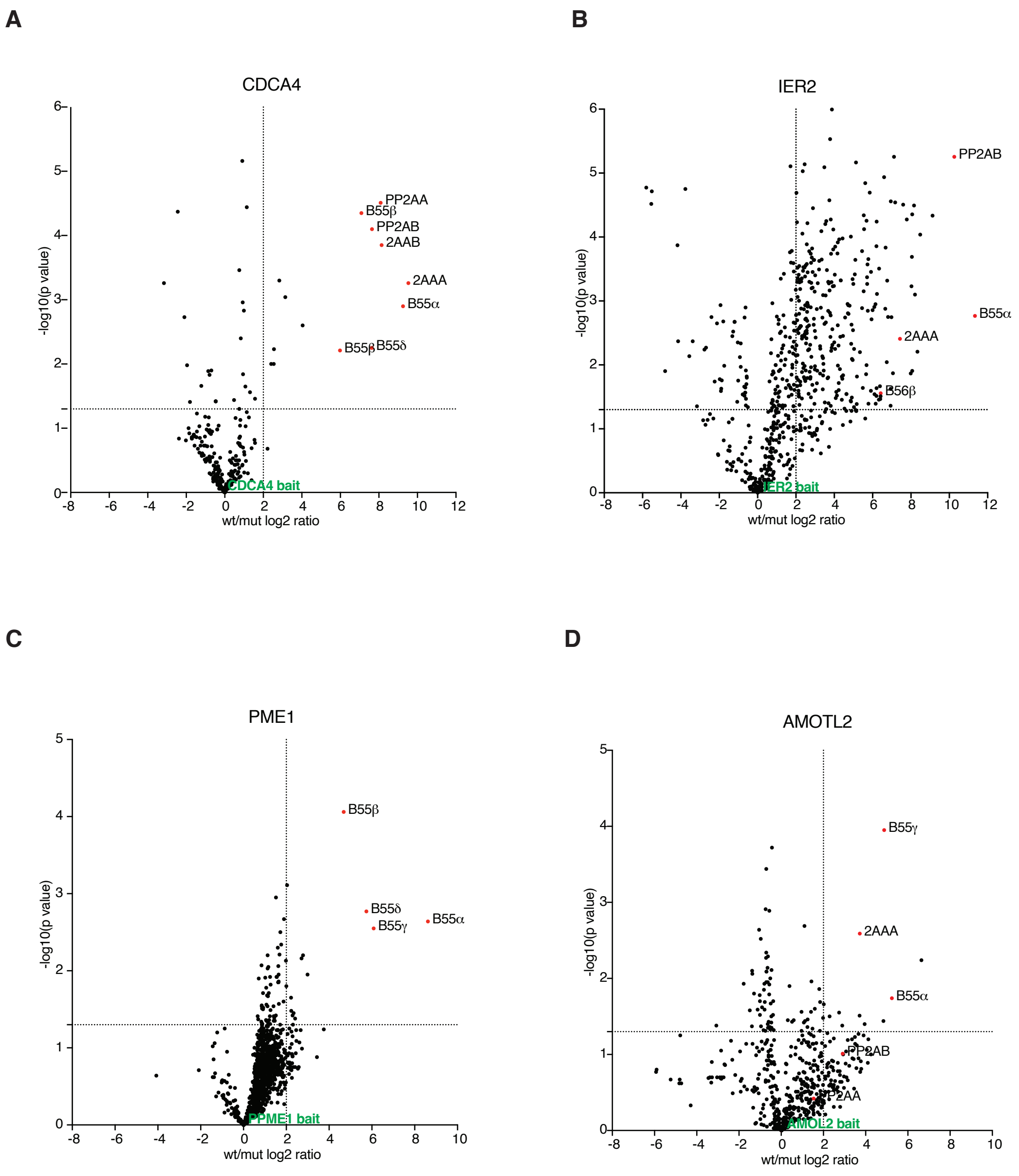


**Supplemental Figure S12**

**A-D)** Volcano plots of mass spectrometry analysis of immunopurifications (n=4) of the indicated proteins comparing wild type to B55 mutant. PP2AA/B: catalytic subunit, 2AAA: scaffold subunit.

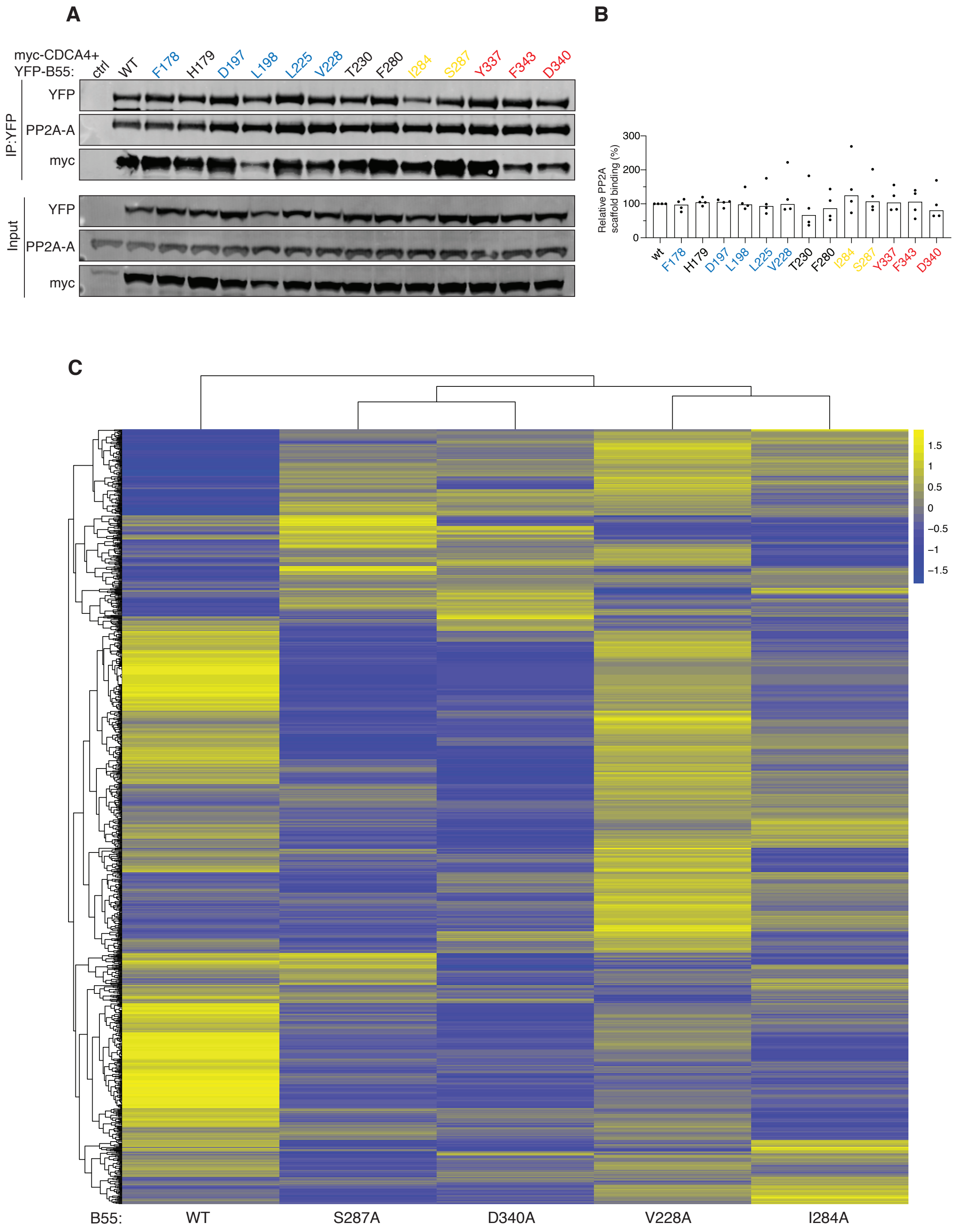


**Supplemental Figure S13**

**A**) Immunoprecipitation of indicated YFP-B55 tagged mutants and binding to myc-CDCA4 determined by western blot **B)** Quantification to relative binding of PP2A scaffold to B55 mutants; median shown, n=4. **C**) HEAT map relating to Figure 2 showing all proteins identified by mass spectrometry in the different B55 variants. Scale bar is log2 and n=4.

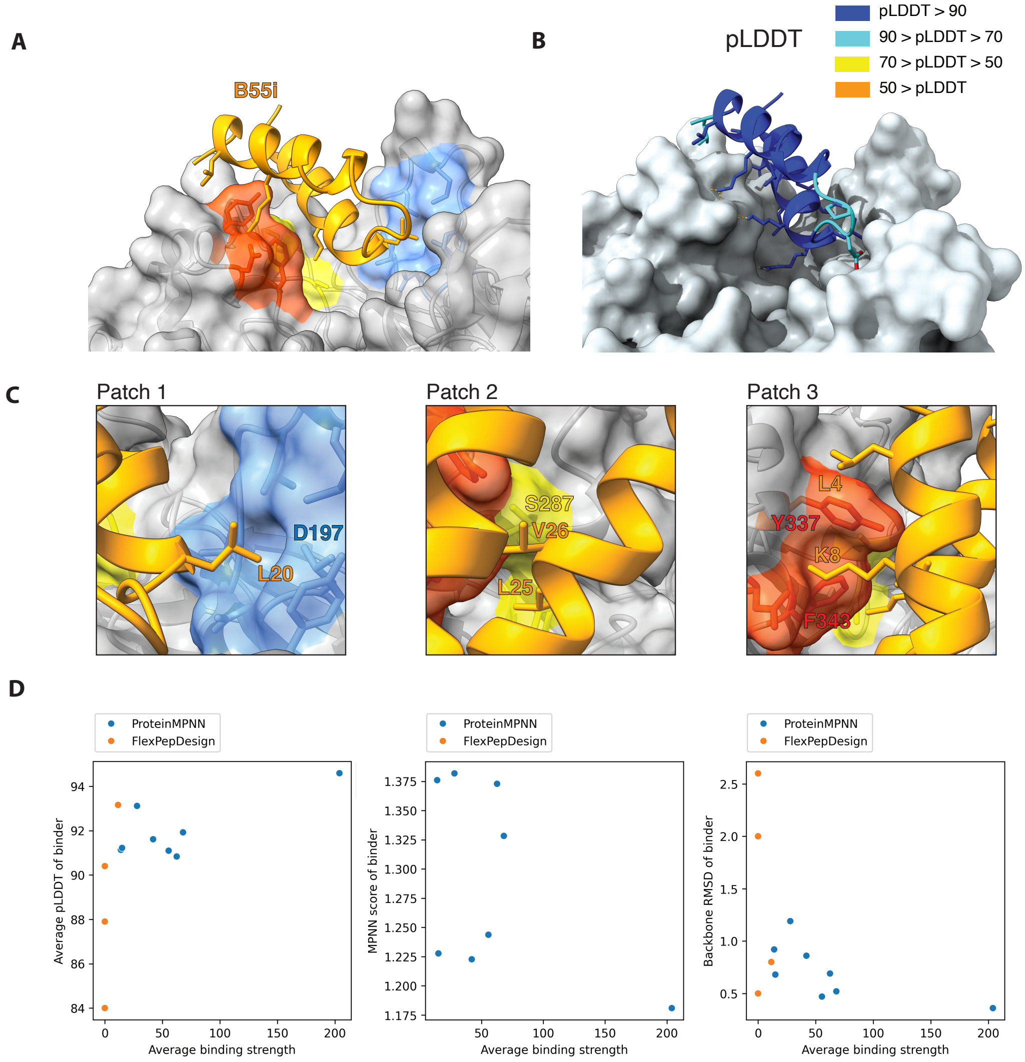


**Supplemental Figure S14**

**A-B**) AF model of B55i bound to B55 and pLDDT score. **C**) Zoom in on interactions of B55i with the different patches. **D**) Plots of design scores and binding strength based on immunopurifications from HeLa cells of YFP tagged de novo B55 binders.

**D**

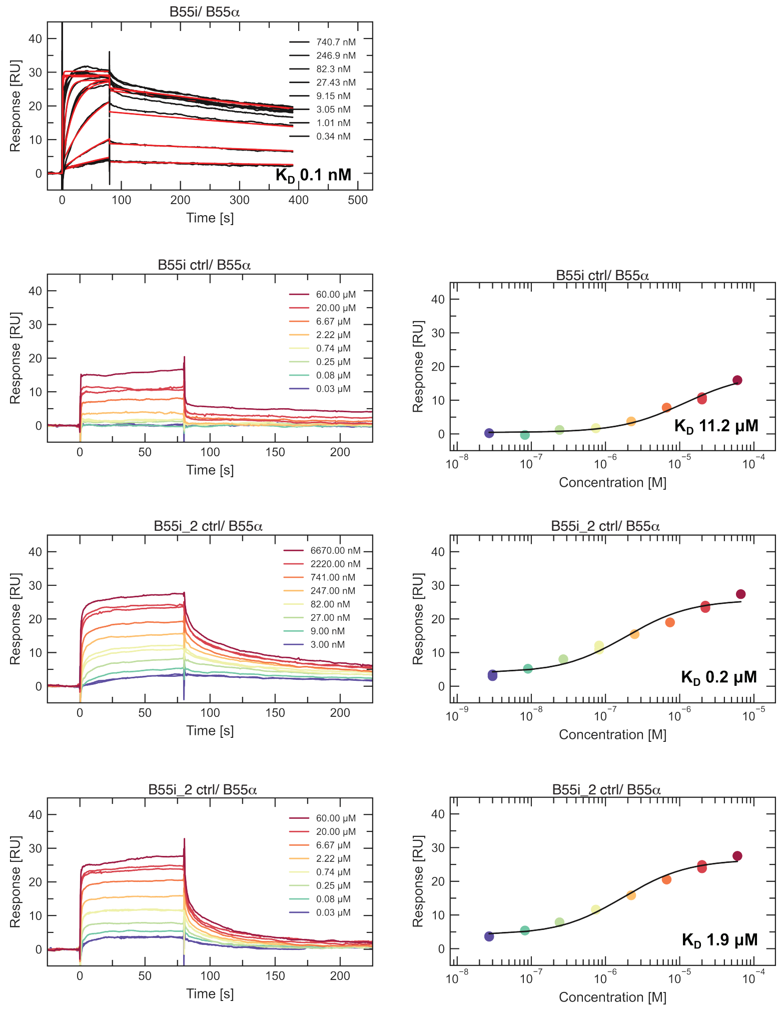


**E**

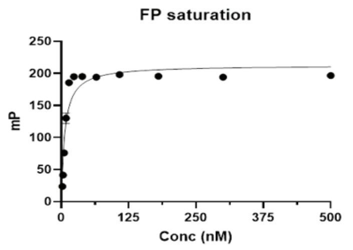


**F**

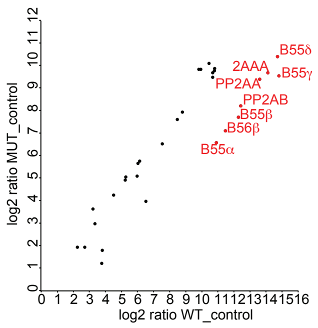


**Supplemental Figure S14-continued**

**D**) SPR data for B55i-1 and B55i-2 peptides binding to B55. **E**) FP measurements for B55i (n=3). **F**) Mass spectrometry analysis of B55i and B55i CTRL based on n=4.

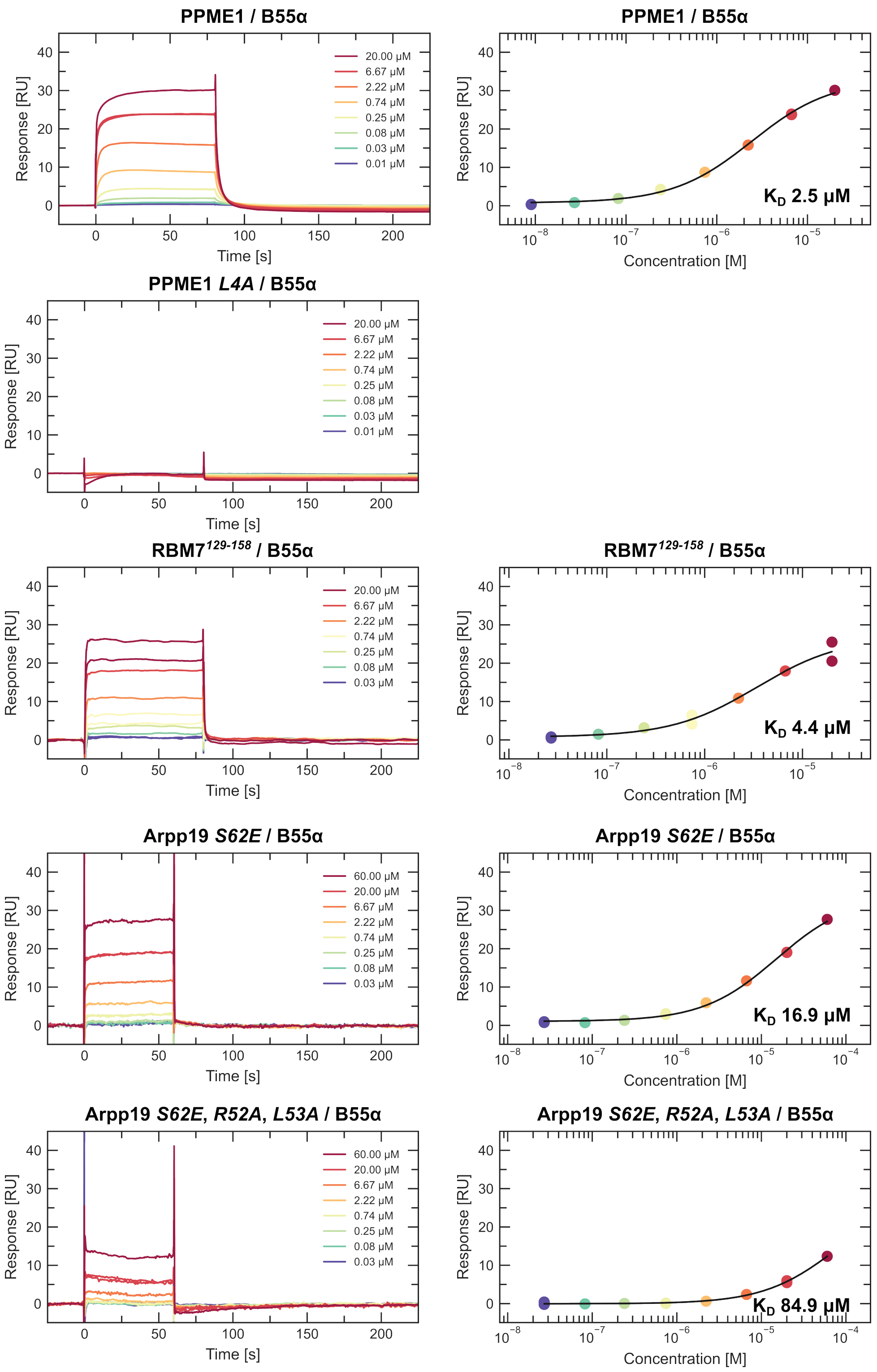


**Supplemental Figure S15**

SPR data for the indicated proteins binding to B55.

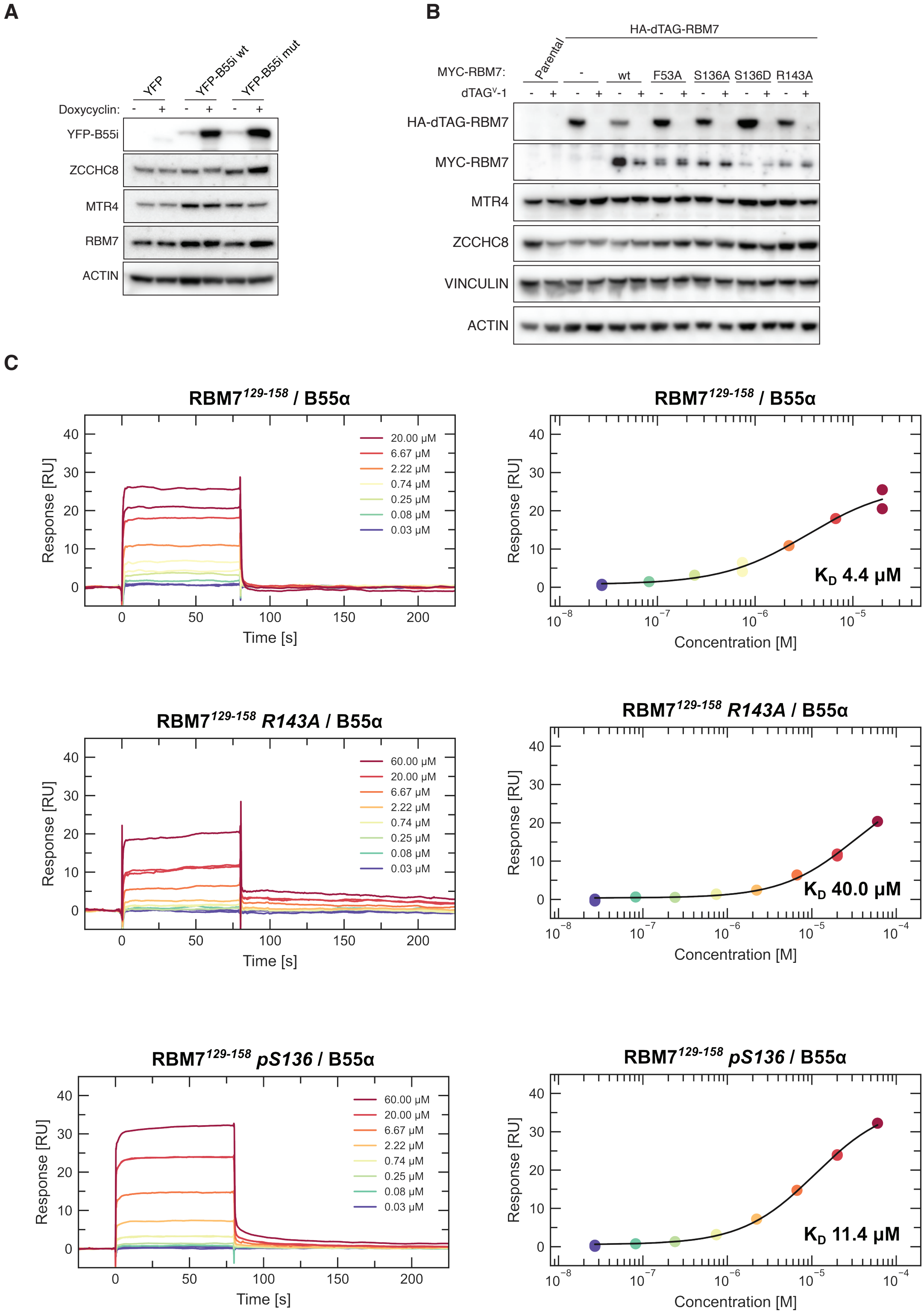


**Supplemental Figure S16**

**A)** Western blots of B55i expression from inducible HeLa cells. **B**) Removal of HA-dTAG-RBM7 with dTAG-1 and complementation with myc-RBM7 variants. **C**) SPR data for the indicated RBM7 peptides binding to B55.
