## Supplemental Table 1 for "Substrate recognition principles for the PP2A-B55 protein phosphatase"

### af2_scores

| pdb | int_plddt | iPAE |
|---|---|---|
| b55_A1X283-SPD2B_HUMAN_unrelaxed_rank_001_alphafold2_multimer_v3_model_1_seed_000.pdb | 73.73 | 9.26 |
| b55_A2A3K4-PTPC1_HUMAN_unrelaxed_rank_001_alphafold2_multimer_v3_model_1_seed_000.pdb | 0.0 | 24.55 |
| b55_A3KMH1-VWA8_HUMAN_unrelaxed_rank_001_alphafold2_multimer_v3_model_5_seed_000.pdb | 65.31 | 26.28 |
| b55_A3KN83-SBNO1_HUMAN_unrelaxed_rank_001_alphafold2_multimer_v3_model_4_seed_000.pdb | 0.0 | 22.28 |
| b55_A7KAX9-RHG32_HUMAN_unrelaxed_rank_001_alphafold2_multimer_v3_model_3_seed_000.pdb | 58.69 | 15.41 |
| b55_A8MPP1-D11L8_HUMAN_unrelaxed_rank_001_alphafold2_multimer_v3_model_4_seed_000.pdb | 72.52 | 27.19 |
| b55_O00165-HAX1_HUMAN_unrelaxed_rank_001_alphafold2_multimer_v3_model_1_seed_000.pdb | 0 | 30 |
| b55_O00311-CDC7_HUMAN_unrelaxed_rank_004_alphafold2_multimer_v3_model_2_seed_000.pdb | 62.44 | 22.7 |
| b55_O00505-IMA4_HUMAN_unrelaxed_rank_003_alphafold2_multimer_v3_model_2_seed_000.pdb | 0.0 | 25.27 |
| b55_O14530-TXND9_HUMAN_unrelaxed_rank_002_alphafold2_multimer_v3_model_3_seed_000.pdb | 73.14 | 26.31 |
| b55_O14734-ACOT8_HUMAN_unrelaxed_rank_003_alphafold2_multimer_v3_model_1_seed_000.pdb | 0 | 18.86 |
| b55_O14737-PDCD5_HUMAN_unrelaxed_rank_001_alphafold2_multimer_v3_model_4_seed_000.pdb | 84.41 | 4.27 |
| b55_O14795-UN13B_HUMAN_unrelaxed_rank_001_alphafold2_multimer_v3_model_2_seed_000.pdb | 65.47 | 10.72 |
| b55_O14907-TX1B3_HUMAN_unrelaxed_rank_001_alphafold2_multimer_v3_model_3_seed_000.pdb | 71.1 | 22.62 |
| b55_O15061-SYNEM_HUMAN_unrelaxed_rank_001_alphafold2_multimer_v3_model_1_seed_000.pdb | 0 | 30 |
| b55_O43166-SI1L1_HUMAN_unrelaxed_rank_001_alphafold2_multimer_v3_model_1_seed_000.pdb | 69.19 | 21.58 |
| b55_O43593-HAIR_HUMAN_unrelaxed_rank_001_alphafold2_multimer_v3_model_4_seed_000.pdb | 65.95 | 8.05 |
| b55_O43768-ENSA_HUMAN_unrelaxed_rank_002_alphafold2_multimer_v3_model_5_seed_000.pdb | 77.41 | 5.06 |
| b55_O43819-SCO2_HUMAN_unrelaxed_rank_001_alphafold2_multimer_v3_model_2_seed_000.pdb | 0.0 | 14.35 |
| b55_O60343-TBCD4_HUMAN_unrelaxed_rank_001_alphafold2_multimer_v3_model_1_seed_000.pdb | 64.14 | 14.22 |
| b55_O60548-FOXD2_HUMAN_unrelaxed_rank_002_alphafold2_multimer_v3_model_5_seed_000.pdb | 79.68 | 6.23 |
| b55_O60610-DIAP1_HUMAN_unrelaxed_rank_001_alphafold2_multimer_v3_model_2_seed_000.pdb | 0 | 30 |
| b55_O60884-DNJA2_HUMAN_unrelaxed_rank_001_alphafold2_multimer_v3_model_1_seed_000.pdb | 0 | 30 |
| b55_O60925-PFD1_HUMAN_unrelaxed_rank_003_alphafold2_multimer_v3_model_4_seed_000.pdb | 0 | 12.97 |
| b55_O75190-DNJB6_HUMAN_unrelaxed_rank_005_alphafold2_multimer_v3_model_2_seed_000.pdb | 64.31 | 23.16 |
| b55_O75376-NCOR1_HUMAN_unrelaxed_alphafold2_multimer_v3_model_2_seed_000.pdb | 0 | 12.58 |
| b55_O75665-OFD1_HUMAN_unrelaxed_rank_001_alphafold2_multimer_v3_model_1_seed_000.pdb | 0 | 30 |
| b55_O75792-RNH2A_HUMAN_unrelaxed_rank_001_alphafold2_multimer_v3_model_5_seed_000.pdb | 0 | 30 |
| b55_O94763-RMP_HUMAN_unrelaxed_rank_004_alphafold2_multimer_v3_model_1_seed_000.pdb | 0.0 | 26.75 |
| b55_O95071-UBR5_HUMAN_unrelaxed_alphafold2_multimer_v3_model_1_seed_000.pdb | 0 | 30 |
| b55_O95263-PDE8B_HUMAN_unrelaxed_rank_004_alphafold2_multimer_v3_model_2_seed_000.pdb | 77.47 | 26.86 |
| b55_O95816-BAG2_HUMAN_unrelaxed_rank_002_alphafold2_multimer_v3_model_1_seed_000.pdb | 63.8 | 30 |
| b55_O95817-BAG3_HUMAN_unrelaxed_rank_001_alphafold2_multimer_v3_model_4_seed_000.pdb | 61.61 | 25.19 |
| b55_P01215-GLHA_HUMAN_unrelaxed_rank_004_alphafold2_multimer_v3_model_3_seed_000.pdb | 0 | 24.62 |
| b55_P04792-HSPB1_HUMAN_unrelaxed_rank_001_alphafold2_multimer_v3_model_2_seed_000.pdb | 0.0 | 7.35 |
| b55_P07900-HS90A_HUMAN_unrelaxed_rank_001_alphafold2_multimer_v3_model_5_seed_000.pdb | 62.79 | 22.92 |
| b55_P08238-HS90B_HUMAN_unrelaxed_rank_001_alphafold2_multimer_v3_model_5_seed_000.pdb | 66.88 | 22.44 |
| b55_P0CG12-DERPC_HUMAN_unrelaxed_rank_001_alphafold2_multimer_v3_model_5_seed_000.pdb | 82.4 | 3.89 |

| |
|---|
| b55_P11498-PYC_HUMAN_unrelaxed_rank_003_alphafold2_multimer_v3_model_4_seed_000.pdb 72.93 26.23 |
| b55_P12036-NFH_HUMAN_unrelaxed_rank_001_alphafold2_multimer_v3_model_2_seed_000.pdb 0 30 |
| b55_P12270-TPR_HUMAN_unrelaxed_rank_001_alphafold2_multimer_v3_model_1_seed_000.pdb 0 30 |
| b55_P13051-UNG_HUMAN_unrelaxed_rank_002_alphafold2_multimer_v3_model_5_seed_000.pdb 79.4 25.72 |
| b55_P17987-TCPA_HUMAN_unrelaxed_rank_001_alphafold2_multimer_v3_model_4_seed_000.pdb 0 30 |
| b55_P19387-RPB3_HUMAN_unrelaxed_rank_001_alphafold2_multimer_v3_model_5_seed_000.pdb 0 30 |
| b55_P23458-JAK1_HUMAN_unrelaxed_rank_003_alphafold2_multimer_v3_model_4_seed_000.pdb 67.14 27.44 |
| b55_P27540-ARNT_HUMAN_unrelaxed_rank_001_alphafold2_multimer_v3_model_2_seed_000.pdb 57.84 15.93 |
| b55_P27708-PYR1_HUMAN_unrelaxed_rank_003_alphafold2_multimer_v3_model_4_seed_000.pdb 75.68 27.66 |
| b55_P29218-IMPA1_HUMAN_unrelaxed_rank_001_alphafold2_multimer_v3_model_3_seed_000.pdb 72.58 26.89 |
| b55_P30876-RPB2_HUMAN_unrelaxed_rank_001_alphafold2_multimer_v3_model_2_seed_000.pdb 0 30 |
| b55_P31689-DNJA1_HUMAN_unrelaxed_rank_001_alphafold2_multimer_v3_model_3_seed_000.pdb 66.39 26.02 |
| b55_P31948-STIP1_HUMAN_unrelaxed_rank_002_alphafold2_multimer_v3_model_1_seed_000.pdb 63.17 22.95 |
| b55_P32926-DSG3_HUMAN_unrelaxed_rank_001_alphafold2_multimer_v3_model_4_seed_000.pdb 84.95 5.93 |
| b55_P36873-PP1G_HUMAN_unrelaxed_rank_002_alphafold2_multimer_v3_model_1_seed_000.pdb 78.52 19.84 |
| b55_P40227-TCPZ_HUMAN_unrelaxed_rank_002_alphafold2_multimer_v3_model_3_seed_000.pdb 0 26.66 |
| b55_P42345-MTOR_HUMAN_unrelaxed_alphafold2_multimer_v3_model_1_seed_000.pdb 83.96 27.75 |
| b55_P48552-NRIP1_HUMAN_unrelaxed_rank_001_alphafold2_multimer_v3_model_4_seed_000.pdb 53.92 12.51 |
| b55_P48643-TCPE_HUMAN_unrelaxed_rank_001_alphafold2_multimer_v3_model_1_seed_000.pdb 0 16.91 |
| b55_P48681-NEST_HUMAN_unrelaxed_rank_001_alphafold2_multimer_v3_model_4_seed_000.pdb 0 30 |
| b55_P49368-TCPG_HUMAN_unrelaxed_rank_001_alphafold2_multimer_v3_model_2_seed_000.pdb 0 30 |
| b55_P50990-TCPQ_HUMAN_unrelaxed_rank_001_alphafold2_multimer_v3_model_2_seed_000.pdb 0.0 25.45 |
| b55_P50991-TCPD_HUMAN_unrelaxed_rank_004_alphafold2_multimer_v3_model_4_seed_000.pdb 0 26.36 |
| b55_P53041-PPP5_HUMAN_unrelaxed_rank_001_alphafold2_multimer_v3_model_1_seed_000.pdb 68.24 24.08 |
| b55_P53634-CATC_HUMAN_unrelaxed_rank_002_alphafold2_multimer_v3_model_1_seed_000.pdb 66.76 26.22 |
| b55_P53814-SMTN_HUMAN_unrelaxed_rank_002_alphafold2_multimer_v3_model_5_seed_000.pdb 70.95 20.64 |
| b55_P55789-ALR_HUMAN_unrelaxed_rank_001_alphafold2_multimer_v3_model_2_seed_000.pdb 0 11.52 |
| b55_P56211-ARP19_HUMAN_unrelaxed_rank_005_alphafold2_multimer_v3_model_4_seed_000.pdb 77.56 4.71 |
| b55_P56524-HDAC4_HUMAN_unrelaxed_rank_001_alphafold2_multimer_v3_model_1_seed_000.pdb 54.55 12.74 |
| b55_P60510-PP4C_HUMAN_unrelaxed_rank_001_alphafold2_multimer_v3_model_3_seed_000.pdb 78.39 8.01 |
| b55_P61758-PFD3_HUMAN_unrelaxed_rank_004_alphafold2_multimer_v3_model_4_seed_000.pdb 65.94 30 |
| b55_P62136-PP1A_HUMAN_unrelaxed_rank_002_alphafold2_multimer_v3_model_1_seed_000.pdb 79.75 22.11 |
| b55_P62140-PP1B_HUMAN_unrelaxed_rank_001_alphafold2_multimer_v3_model_5_seed_000.pdb 0.0 13.51 |
| b55_P63151-2ABA_HUMAN_unrelaxed_rank_002_alphafold2_multimer_v3_model_3_seed_000.pdb 72.77 23.09 |
| b55_P78371-TCPB_HUMAN_unrelaxed_rank_003_alphafold2_multimer_v3_model_4_seed_000.pdb 65.46 26.69 |
| b55_P86791-CCZ1_HUMAN_unrelaxed_rank_003_alphafold2_multimer_v3_model_4_seed_000.pdb 0.0 16.38 |
| b55_Q00005-2ABB_HUMAN_unrelaxed_rank_002_alphafold2_multimer_v3_model_1_seed_000.pdb 75.07 21.14 |
| b55_Q00587-BORG5_HUMAN_unrelaxed_rank_001_alphafold2_multimer_v3_model_4_seed_000.pdb 70.91 7.94 |
| b55_Q05209-PTN12_HUMAN_unrelaxed_rank_002_alphafold2_multimer_v3_model_3_seed_000.pdb 76.67 20.7 |
| b55_Q08209-PP2BA_HUMAN_unrelaxed_rank_003_alphafold2_multimer_v3_model_1_seed_000.pdb 0.0 26.68 |

| |
|---|
| b55_Q08AF3-SLFN5_HUMAN_unrelaxed_rank_002_alphafold2_multimer_v3_model_1_seed_000.pdb 69.07 27.17 |
| b55_Q0VG06-FP100_HUMAN_unrelaxed_rank_002_alphafold2_multimer_v3_model_2_seed_000.pdb 68.07 26.7 |
| b55_Q12815-TROAP_HUMAN_unrelaxed_rank_001_alphafold2_multimer_v3_model_2_seed_000.pdb 73.87 5.38 |
| b55_Q12851-M4K2_HUMAN_unrelaxed_rank_002_alphafold2_multimer_v3_model_5_seed_000.pdb 71.28 25.25 |
| b55_Q12948-FOXC1_HUMAN_unrelaxed_rank_001_alphafold2_multimer_v3_model_3_seed_000.pdb 84.19 4.39 |
| b55_Q13015-AF1Q_HUMAN_unrelaxed_rank_001_alphafold2_multimer_v3_model_3_seed_000.pdb 0 30 |
| b55_Q13049-TRI32_HUMAN_unrelaxed_rank_004_alphafold2_multimer_v3_model_4_seed_000.pdb 67.48 24.38 |
| b55_Q13322-GRB10_HUMAN_unrelaxed_rank_002_alphafold2_multimer_v3_model_1_seed_000.pdb 68.71 26.3 |
| b55_Q13371-PHLP_HUMAN_unrelaxed_rank_001_alphafold2_multimer_v3_model_3_seed_000.pdb 75.4 7.03 |
| b55_Q13451-FKBP5_HUMAN_unrelaxed_rank_002_alphafold2_multimer_v3_model_2_seed_000.pdb 74.06 19.1 |
| b55_Q13523-PRP4B_HUMAN_unrelaxed_rank_001_alphafold2_multimer_v3_model_1_seed_000.pdb 60.56 12.73 |
| b55_Q13671-RIN1_HUMAN_unrelaxed_rank_001_alphafold2_multimer_v3_model_4_seed_000.pdb 72.99 5.86 |
| b55_Q14140-SRTD2_HUMAN_unrelaxed_rank_003_alphafold2_multimer_v3_model_4_seed_000.pdb 92.53 2.16 |
| b55_Q14678-KANK1_HUMAN_unrelaxed_rank_001_alphafold2_multimer_v3_model_3_seed_000.pdb 0 9.57 |
| b55_Q15031-SYLM_HUMAN_unrelaxed_rank_002_alphafold2_multimer_v3_model_3_seed_000.pdb 75.96 26.16 |
| b55_Q15438-CYH1_HUMAN_unrelaxed_rank_001_alphafold2_multimer_v3_model_2_seed_000.pdb 69.31 11.83 |
| b55_Q15468-STIL_HUMAN_unrelaxed_rank_002_alphafold2_multimer_v3_model_4_seed_000.pdb 0.0 13.74 |
| b55_Q15773-MLF2_HUMAN_unrelaxed_rank_001_alphafold2_multimer_v3_model_2_seed_000.pdb 68.78 9.06 |
| b55_Q16342-PDCD2_HUMAN_unrelaxed_rank_004_alphafold2_multimer_v3_model_5_seed_000.pdb 82.2 24.47 |
| b55_Q16563-SYPL1_HUMAN_unrelaxed_rank_001_alphafold2_multimer_v3_model_2_seed_000.pdb 59.95 12.11 |
| b55_Q16584-M3K11_HUMAN_unrelaxed_rank_001_alphafold2_multimer_v3_model_5_seed_000.pdb 66.74 24.78 |
| b55_Q1MSJ5-CSPP1_HUMAN_unrelaxed_rank_001_alphafold2_multimer_v3_model_2_seed_000.pdb 77.31 5.68 |
| b55_Q2KHR3-QSER1_HUMAN_unrelaxed_rank_002_alphafold2_multimer_v3_model_1_seed_000.pdb 59.27 23.66 |
| b55_Q2M3G4-SHRM1_HUMAN_unrelaxed_rank_003_alphafold2_multimer_v3_model_3_seed_000.pdb 73.59 15.77 |
| b55_Q32MH5-ATOSA_HUMAN_unrelaxed_rank_002_alphafold2_multimer_v3_model_5_seed_000.pdb 89.82 3.27 |
| b55_Q3YEC7-RABL6_HUMAN_unrelaxed_rank_001_alphafold2_multimer_v3_model_4_seed_000.pdb 0.0 17.72 |
| b55_Q53ET0-CRTC2_HUMAN_unrelaxed_rank_001_alphafold2_multimer_v3_model_5_seed_000.pdb 0 30 |
| b55_Q58FF8-H90B2_HUMAN_unrelaxed_rank_002_alphafold2_multimer_v3_model_2_seed_000.pdb 0.0 22.53 |
| b55_Q58WW2-DCAF6_HUMAN_unrelaxed_rank_001_alphafold2_multimer_v3_model_5_seed_000.pdb 0.0 26.14 |
| b55_Q5M775-CYTSB_HUMAN_unrelaxed_rank_001_alphafold2_multimer_v3_model_4_seed_000.pdb 71.56 9.81 |
| b55_Q5T447-HECD3_HUMAN_unrelaxed_rank_002_alphafold2_multimer_v3_model_2_seed_000.pdb 71.27 27.62 |
| b55_Q5T5U3-RHG21_HUMAN_unrelaxed_rank_001_alphafold2_multimer_v3_model_5_seed_000.pdb 0.0 8.52 |
| b55_Q5T953-IER5L_HUMAN_unrelaxed_rank_001_alphafold2_multimer_v3_model_4_seed_000.pdb 90.5 2.03 |
| b55_Q5T9S5-CCD18_HUMAN_unrelaxed_rank_001_alphafold2_multimer_v3_model_4_seed_000.pdb 0 30 |
| b55_Q5VT52-RPRD2_HUMAN_unrelaxed_rank_001_alphafold2_multimer_v3_model_4_seed_000.pdb 0.0 8.1 |
| b55_Q5VY09-IER5_HUMAN_unrelaxed_rank_002_alphafold2_multimer_v3_model_2_seed_000.pdb 90.91 2.43 |
| b55_Q63ZY3-KANK2_HUMAN_unrelaxed_rank_003_alphafold2_multimer_v3_model_4_seed_000.pdb 56.69 13.2 |
| b55_Q66LE6-2ABD_HUMAN_unrelaxed_rank_001_alphafold2_multimer_v3_model_1_seed_000.pdb 74.6 21.59 |
| b55_Q6BDS2-BLT3A_HUMAN_unrelaxed_rank_005_alphafold2_multimer_v3_model_4_seed_000.pdb 80.91 26.42 |
| b55_Q6NYC8-PPR18_HUMAN_unrelaxed_rank_001_alphafold2_multimer_v3_model_4_seed_000.pdb 0 30 |

| |
|---|
| b55_Q6NZY4-ZCHC8_HUMAN_unrelaxed_rank_001_alphafold2_multimer_v3_model_1_seed_000.pdb 0 30 |
| b55_Q6P3X3-TTC27_HUMAN_unrelaxed_rank_003_alphafold2_multimer_v3_model_3_seed_000.pdb 74.7 28.36 |
| b55_Q6P996-PDXD1_HUMAN_unrelaxed_rank_002_alphafold2_multimer_v3_model_2_seed_000.pdb 80.79 27.48 |
| b55_Q6PID6-TTC33_HUMAN_unrelaxed_rank_001_alphafold2_multimer_v3_model_1_seed_000.pdb 73.23 25.44 |
| b55_Q6UUV7-CRTC3_HUMAN_unrelaxed_rank_001_alphafold2_multimer_v3_model_5_seed_000.pdb 0 30 |
| b55_Q70CQ4-UBP31_HUMAN_unrelaxed_rank_002_alphafold2_multimer_v3_model_4_seed_000.pdb 0.0 10.02 |
| b55_Q70EL4-UBP43_HUMAN_unrelaxed_rank_001_alphafold2_multimer_v3_model_5_seed_000.pdb 63.92 9.12 |
| b55_Q7RTV0-PHF5A_HUMAN_unrelaxed_rank_001_alphafold2_multimer_v3_model_2_seed_000.pdb 0.0 20.2 |
| b55_Q7Z401-MYCPP_HUMAN_unrelaxed_rank_003_alphafold2_multimer_v3_model_1_seed_000.pdb 68.42 27.59 |
| b55_Q7Z7K0-COXM1_HUMAN_unrelaxed_rank_002_alphafold2_multimer_v3_model_4_seed_000.pdb 59.97 22.11 |
| b55_Q86SQ0-PHLB2_HUMAN_unrelaxed_rank_003_alphafold2_multimer_v3_model_4_seed_000.pdb 0.0 25.94 |
| b55_Q86TC9-MYPN_HUMAN_unrelaxed_rank_003_alphafold2_multimer_v3_model_5_seed_000.pdb 0.0 21.66 |
| b55_Q86U70-LDB1_HUMAN_unrelaxed_rank_001_alphafold2_multimer_v3_model_1_seed_000.pdb 0.0 24.52 |
| b55_Q86UU1-PHLB1_HUMAN_unrelaxed_rank_002_alphafold2_multimer_v3_model_3_seed_000.pdb 73.11 13.88 |
| b55_Q8IY67-RAVR1_HUMAN_unrelaxed_rank_004_alphafold2_multimer_v3_model_2_seed_000.pdb 0.0 23.84 |
| b55_Q8N157-AHI1_HUMAN_unrelaxed_rank_001_alphafold2_multimer_v3_model_3_seed_000.pdb 80.41 13.94 |
| b55_Q8N163-CCAR2_HUMAN_unrelaxed_rank_002_alphafold2_multimer_v3_model_2_seed_000.pdb 0 27.78 |
| b55_Q8N5M4-TTC9C_HUMAN_unrelaxed_rank_001_alphafold2_multimer_v3_model_1_seed_000.pdb 75.21 17.14 |
| b55_Q8N612-FHI1B_HUMAN_unrelaxed_rank_005_alphafold2_multimer_v3_model_2_seed_000.pdb 86.14 28.33 |
| b55_Q8N8E3-CE112_HUMAN_unrelaxed_rank_003_alphafold2_multimer_v3_model_1_seed_000.pdb 67.59 26.52 |
| b55_Q8NDG6-TDRD9_HUMAN_unrelaxed_rank_001_alphafold2_multimer_v3_model_4_seed_000.pdb 75.26 26.09 |
| b55_Q8NHY2-COP1_HUMAN_unrelaxed_rank_001_alphafold2_multimer_v3_model_2_seed_000.pdb 83.5 10.85 |
| b55_Q8TBC3-SHKB1_HUMAN_unrelaxed_rank_002_alphafold2_multimer_v3_model_3_seed_000.pdb 69.2 24.78 |
| b55_Q8TCS8-PNPT1_HUMAN_unrelaxed_rank_002_alphafold2_multimer_v3_model_5_seed_000.pdb 0.0 27.75 |
| b55_Q8TEP8-CE192_HUMAN_unrelaxed_alphafold2_multimer_v3_model_1_seed_000.pdb 68.18 27.75 |
| b55_Q8TEU7-RPGF6_HUMAN_unrelaxed_rank_001_alphafold2_multimer_v3_model_1_seed_000.pdb 0.0 27.08 |
| b55_Q8TF61-FBX41_HUMAN_unrelaxed_rank_001_alphafold2_multimer_v3_model_2_seed_000.pdb 68.55 8.16 |
| b55_Q8WUF5-IASPP_HUMAN_unrelaxed_rank_001_alphafold2_multimer_v3_model_2_seed_000.pdb 60.77 9.71 |
| b55_Q8WUI4-HDAC7_HUMAN_unrelaxed_rank_002_alphafold2_multimer_v3_model_1_seed_000.pdb 69.85 26.27 |
| b55_Q8WUK0-PTPM1_HUMAN_unrelaxed_rank_001_alphafold2_multimer_v3_model_3_seed_000.pdb 82.57 4.2 |
| b55_Q8WUM9-S20A1_HUMAN_unrelaxed_rank_001_alphafold2_multimer_v3_model_5_seed_000.pdb 79.37 3.87 |
| b55_Q8WV74-NUDT8_HUMAN_unrelaxed_rank_001_alphafold2_multimer_v3_model_5_seed_000.pdb 64.39 12.93 |
| b55_Q8WW01-SEN15_HUMAN_unrelaxed_rank_002_alphafold2_multimer_v3_model_1_seed_000.pdb 68.47 20.39 |
| b55_Q8WXG6-MADD_HUMAN_unrelaxed_rank_001_alphafold2_multimer_v3_model_1_seed_000.pdb 0.0 28.11 |
| b55_Q92526-TCPW_HUMAN_unrelaxed_rank_001_alphafold2_multimer_v3_model_4_seed_000.pdb 0 30 |
| b55_Q92540-SMG7_HUMAN_unrelaxed_rank_002_alphafold2_multimer_v3_model_5_seed_000.pdb 64.35 15.07 |
| b55_Q92771-DDX12_HUMAN_unrelaxed_rank_002_alphafold2_multimer_v3_model_4_seed_000.pdb 0.0 27.2 |
| b55_Q96A73-P33MX_HUMAN_unrelaxed_rank_001_alphafold2_multimer_v3_model_3_seed_000.pdb 62.11 10.34 |
| b55_Q96AT1-K1143_HUMAN_unrelaxed_rank_001_alphafold2_multimer_v3_model_4_seed_000.pdb 0 13.99 |
| b55_Q96D53-COQ8B_HUMAN_unrelaxed_rank_003_alphafold2_multimer_v3_model_3_seed_000.pdb 0.0 23.81 |

| |
|---|
| b55_Q96D71-REPS1_HUMAN_unrelaxed_rank_002_alphafold2_multimer_v3_model_1_seed_000.pdb 0 13.81 |
| b55_Q96DF8-ESS2_HUMAN_unrelaxed_rank_001_alphafold2_multimer_v3_model_1_seed_000.pdb 63.71 10.33 |
| b55_Q96E09-PBIR1_HUMAN_unrelaxed_rank_001_alphafold2_multimer_v3_model_4_seed_000.pdb 82.4 2.99 |
| b55_Q96FC9-DDX11_HUMAN_unrelaxed_rank_002_alphafold2_multimer_v3_model_4_seed_000.pdb 75.57 26.98 |
| b55_Q96HB5-CC120_HUMAN_unrelaxed_rank_003_alphafold2_multimer_v3_model_3_seed_000.pdb 0 22.55 |
| b55_Q96II8-LRCH3_HUMAN_unrelaxed_rank_001_alphafold2_multimer_v3_model_2_seed_000.pdb 72.13 9.43 |
| b55_Q96IU2-ZBED3_HUMAN_unrelaxed_rank_003_alphafold2_multimer_v3_model_3_seed_000.pdb 0 25.5 |
| b55_Q96N67-DOCK7_HUMAN_unrelaxed_rank_001_alphafold2_multimer_v3_model_1_seed_000.pdb 0 30 |
| b55_Q96PE3-INP4A_HUMAN_unrelaxed_rank_001_alphafold2_multimer_v3_model_4_seed_000.pdb 82.04 25.95 |
| b55_Q96QU8-XPO6_HUMAN_unrelaxed_rank_003_alphafold2_multimer_v3_model_2_seed_000.pdb 76.56 27.92 |
| b55_Q96RT1-ERBIN_HUMAN_unrelaxed_rank_001_alphafold2_multimer_v3_model_4_seed_000.pdb 0 26.28 |
| b55_Q99471-PFD5_HUMAN_unrelaxed_rank_001_alphafold2_multimer_v3_model_5_seed_000.pdb 0 30 |
| b55_Q99504-EYA3_HUMAN_unrelaxed_rank_005_alphafold2_multimer_v3_model_4_seed_000.pdb 76.04 25.12 |
| b55_Q99550-MPP9_HUMAN_unrelaxed_rank_002_alphafold2_multimer_v3_model_1_seed_000.pdb 59.99 11.63 |
| b55_Q99832-TCPH_HUMAN_unrelaxed_rank_003_alphafold2_multimer_v3_model_3_seed_000.pdb 83.41 28.53 |
| b55_Q99933-BAG1_HUMAN_unrelaxed_rank_002_alphafold2_multimer_v3_model_5_seed_000.pdb 65.63 20.67 |
| b55_Q99958-FOXC2_HUMAN_unrelaxed_rank_003_alphafold2_multimer_v3_model_4_seed_000.pdb 75.17 8.7 |
| b55_Q99959-PKP2_HUMAN_unrelaxed_rank_004_alphafold2_multimer_v3_model_3_seed_000.pdb 77.97 25.11 |
| b55_Q9BQ95-ECSIT_HUMAN_unrelaxed_rank_003_alphafold2_multimer_v3_model_4_seed_000.pdb 73.54 25.25 |
| b55_Q9BRK5-CAB45_HUMAN_unrelaxed_rank_002_alphafold2_multimer_v3_model_4_seed_000.pdb 71.53 25.22 |
| b55_Q9BRQ8-FSP1_HUMAN_unrelaxed_rank_001_alphafold2_multimer_v3_model_4_seed_000.pdb 96.89 2.41 |
| b55_Q9BSJ6-PIMRE_HUMAN_unrelaxed_rank_001_alphafold2_multimer_v3_model_3_seed_000.pdb 0 9.49 |
| b55_Q9BSV6-SEN34_HUMAN_unrelaxed_rank_001_alphafold2_multimer_v3_model_2_seed_000.pdb 0 30 |
| b55_Q9BTL4-IER2_HUMAN_unrelaxed_rank_002_alphafold2_multimer_v3_model_2_seed_000.pdb 89.46 2.4 |
| b55_Q9BU02-THTPA_HUMAN_unrelaxed_rank_001_alphafold2_multimer_v3_model_3_seed_000.pdb 69.75 21.19 |
| b55_Q9BU61-NDUF3_HUMAN_unrelaxed_rank_002_alphafold2_multimer_v3_model_1_seed_000.pdb 0 24.95 |
| b55_Q9BUP3-HTAI2_HUMAN_unrelaxed_rank_003_alphafold2_multimer_v3_model_5_seed_000.pdb 62.21 18.39 |
| b55_Q9BW92-SYTM_HUMAN_unrelaxed_rank_001_alphafold2_multimer_v3_model_5_seed_000.pdb 77.53 23.48 |
| b55_Q9BWT7-CAR10_HUMAN_unrelaxed_rank_001_alphafold2_multimer_v3_model_4_seed_000.pdb 59.56 19.12 |
| b55_Q9BXF6-RFIP5_HUMAN_unrelaxed_rank_003_alphafold2_multimer_v3_model_2_seed_000.pdb 55.96 17.81 |
| b55_Q9BXL8-CDCA4_HUMAN_unrelaxed_rank_002_alphafold2_multimer_v3_model_4_seed_000.pdb 94.28 1.73 |
| b55_Q9BYV8-CEP41_HUMAN_unrelaxed_rank_003_alphafold2_multimer_v3_model_2_seed_000.pdb 0.0 26.64 |
| b55_Q9BZL6-KPCD2_HUMAN_unrelaxed_rank_002_alphafold2_multimer_v3_model_3_seed_000.pdb 64.39 25.06 |
| b55_Q9C0B7-TNG6_HUMAN_unrelaxed_rank_001_alphafold2_multimer_v3_model_2_seed_000.pdb 0 30 |
| b55_Q9H0K1-SIK2_HUMAN_unrelaxed_rank_001_alphafold2_multimer_v3_model_5_seed_000.pdb 82.48 3.58 |
| b55_Q9H0R5-GBP3_HUMAN_unrelaxed_rank_002_alphafold2_multimer_v3_model_1_seed_000.pdb 0.0 26.39 |
| b55_Q9H1H9-KI13A_HUMAN_unrelaxed_rank_005_alphafold2_multimer_v3_model_4_seed_000.pdb 66.54 26.64 |
| b55_Q9H2J4-PDCL3_HUMAN_unrelaxed_rank_004_alphafold2_multimer_v3_model_3_seed_000.pdb 0.0 23.97 |
| b55_Q9H4G0-E41L1_HUMAN_unrelaxed_rank_001_alphafold2_multimer_v3_model_5_seed_000.pdb 84.2 5.02 |
| b55_Q9H4M9-EHD1_HUMAN_unrelaxed_rank_002_alphafold2_multimer_v3_model_1_seed_000.pdb 69.63 19.06 |

| |
|---|
| b55_Q9H5N1-RABE2_HUMAN_unrelaxed_rank_001_alphafold2_multimer_v3_model_3_seed_000.pdb 61.69 17.72 |
| b55_Q9H5Z1-DHX35_HUMAN_unrelaxed_rank_001_alphafold2_multimer_v3_model_5_seed_000.pdb 76.47 12.42 |
| b55_Q9H845-ACAD9_HUMAN_unrelaxed_rank_001_alphafold2_multimer_v3_model_5_seed_000.pdb 0.0 18.16 |
| b55_Q9H910-JUPI2_HUMAN_unrelaxed_rank_001_alphafold2_multimer_v3_model_3_seed_000.pdb 0 11.65 |
| b55_Q9HCN8-SDF2L_HUMAN_unrelaxed_rank_001_alphafold2_multimer_v3_model_3_seed_000.pdb 83.72 24.62 |
| b55_Q9NNX1-TUFT1_HUMAN_unrelaxed_rank_001_alphafold2_multimer_v3_model_3_seed_000.pdb 77.25 9.48 |
| b55_Q9NQ35-NRIP3_HUMAN_unrelaxed_rank_001_alphafold2_multimer_v3_model_1_seed_000.pdb 73.09 4.59 |
| b55_Q9NQI0-DDX4_HUMAN_unrelaxed_rank_001_alphafold2_multimer_v3_model_4_seed_000.pdb 82.5 23.61 |
| b55_Q9NQP4-PFD4_HUMAN_unrelaxed_rank_001_alphafold2_multimer_v3_model_3_seed_000.pdb 0 26.58 |
| b55_Q9NQT8-KI13B_HUMAN_unrelaxed_rank_001_alphafold2_multimer_v3_model_4_seed_000.pdb 65.89 9.77 |
| b55_Q9NR46-SHLB2_HUMAN_unrelaxed_rank_001_alphafold2_multimer_v3_model_5_seed_000.pdb 65.73 8.98 |
| b55_Q9NRA8-4ET_HUMAN_unrelaxed_rank_001_alphafold2_multimer_v3_model_4_seed_000.pdb 69.39 7.13 |
| b55_Q9NRL3-STRN4_HUMAN_unrelaxed_rank_001_alphafold2_multimer_v3_model_5_seed_000.pdb 0.0 25.8 |
| b55_Q9NRV9-HEBP1_HUMAN_unrelaxed_rank_001_alphafold2_multimer_v3_model_3_seed_000.pdb 0.0 17.03 |
| b55_Q9NSY1-BMP2K_HUMAN_unrelaxed_rank_001_alphafold2_multimer_v3_model_1_seed_000.pdb 64.61 13.47 |
| b55_Q9NTX5-ECHD1_HUMAN_unrelaxed_rank_005_alphafold2_multimer_v3_model_2_seed_000.pdb 76.48 25.95 |
| b55_Q9NUJ3-T11L1_HUMAN_unrelaxed_rank_002_alphafold2_multimer_v3_model_4_seed_000.pdb 61.26 26.64 |
| b55_Q9NUP9-LIN7C_HUMAN_unrelaxed_rank_001_alphafold2_multimer_v3_model_1_seed_000.pdb 0.0 23.45 |
| b55_Q9NZ71-RTEL1_HUMAN_unrelaxed_rank_001_alphafold2_multimer_v3_model_1_seed_000.pdb 69.82 12.29 |
| b55_Q9NZW5-PALS2_HUMAN_unrelaxed_rank_003_alphafold2_multimer_v3_model_4_seed_000.pdb 0.0 24.89 |
| b55_Q9P0K8-FOXJ2_HUMAN_unrelaxed_rank_001_alphafold2_multimer_v3_model_5_seed_000.pdb 77.34 9.42 |
| b55_Q9P202-WHRN_HUMAN_unrelaxed_rank_001_alphafold2_multimer_v3_model_3_seed_000.pdb 64.86 9.99 |
| b55_Q9P227-RHG23_HUMAN_unrelaxed_rank_002_alphafold2_multimer_v3_model_4_seed_000.pdb 58.69 13.43 |
| b55_Q9P270-SLAI2_HUMAN_unrelaxed_rank_001_alphafold2_multimer_v3_model_4_seed_000.pdb 62.22 14.93 |
| b55_Q9P2I0-CPSF2_HUMAN_unrelaxed_rank_004_alphafold2_multimer_v3_model_1_seed_000.pdb 69.28 27.06 |
| b55_Q9UBI6-GBG12_HUMAN_unrelaxed_rank_001_alphafold2_multimer_v3_model_5_seed_000.pdb 0 30 |
| b55_Q9UDT6-CLIP2_HUMAN_unrelaxed_rank_001_alphafold2_multimer_v3_model_4_seed_000.pdb 80.87 4.48 |
| b55_Q9UDY4-DNJB4_HUMAN_unrelaxed_rank_001_alphafold2_multimer_v3_model_3_seed_000.pdb 73.09 27.41 |
| b55_Q9UGU0-TCF20_HUMAN_unrelaxed_rank_001_alphafold2_multimer_v3_model_4_seed_000.pdb 0.0 9.26 |
| b55_Q9UHB6-LIMA1_HUMAN_unrelaxed_rank_001_alphafold2_multimer_v3_model_4_seed_000.pdb 60.0 11.67 |
| b55_Q9UJY1-HSPB8_HUMAN_unrelaxed_rank_003_alphafold2_multimer_v3_model_2_seed_000.pdb 0 25.83 |
| b55_Q9UKY7-CDV3_HUMAN_unrelaxed_rank_002_alphafold2_multimer_v3_model_2_seed_000.pdb 0 11.74 |
| b55_Q9UL15-BAG5_HUMAN_unrelaxed_rank_001_alphafold2_multimer_v3_model_4_seed_000.pdb 69.34 25.44 |
| b55_Q9UNE7-CHIP_HUMAN_unrelaxed_rank_003_alphafold2_multimer_v3_model_1_seed_000.pdb 66.98 25.16 |
| b55_Q9UPQ0-LIMC1_HUMAN_unrelaxed_rank_001_alphafold2_multimer_v3_model_5_seed_000.pdb 73.4 5.49 |
| b55_Q9UPQ4-TRI35_HUMAN_unrelaxed_rank_001_alphafold2_multimer_v3_model_2_seed_000.pdb 72.63 27.3 |
| b55_Q9UQL6-HDAC5_HUMAN_unrelaxed_rank_002_alphafold2_multimer_v3_model_2_seed_000.pdb 0.0 12.86 |
| b55_Q9Y2H2-SAC2_HUMAN_unrelaxed_rank_001_alphafold2_multimer_v3_model_1_seed_000.pdb 0 14.95 |
| b55_Q9Y2H5-PKHA6_HUMAN_unrelaxed_rank_002_alphafold2_multimer_v3_model_4_seed_000.pdb 68.3 8.59 |
| b55_Q9Y2J4-AMOL2_HUMAN_unrelaxed_rank_001_alphafold2_multimer_v3_model_2_seed_000.pdb 64.32 11.27 |

| |
|---|
| b55_Q9Y2K2-SIK3_HUMAN_unrelaxed_rank_001_alphafold2_multimer_v3_model_1_seed_000.pdb 63.81 10.37 |
| b55_Q9Y2Q3-GSTK1_HUMAN_unrelaxed_rank_001_alphafold2_multimer_v3_model_5_seed_000.pdb 69.74 22.84 |
| b55_Q9Y2Z0-SGT1_HUMAN_unrelaxed_rank_003_alphafold2_multimer_v3_model_2_seed_000.pdb 0.0 25.17 |
| b55_Q9Y371-SHLB1_HUMAN_unrelaxed_rank_001_alphafold2_multimer_v3_model_3_seed_000.pdb 64.44 19.0 |
| b55_Q9Y375-CIA30_HUMAN_unrelaxed_rank_002_alphafold2_multimer_v3_model_2_seed_000.pdb 62.51 16.86 |
| b55_Q9Y3X0-CCDC9_HUMAN_unrelaxed_rank_003_alphafold2_multimer_v3_model_5_seed_000.pdb 0 21.68 |
| b55_Q9Y446-PKP3_HUMAN_unrelaxed_rank_001_alphafold2_multimer_v3_model_1_seed_000.pdb 0.0 11.04 |
| b55_Q9Y4R8-TELO2_HUMAN_unrelaxed_rank_001_alphafold2_multimer_v3_model_4_seed_000.pdb 0 30 |
| b55_Q9Y570-PPME1_HUMAN_unrelaxed_rank_002_alphafold2_multimer_v3_model_5_seed_000.pdb 75.52 5.41 |
| b55_Q9Y580-RBM7_HUMAN_unrelaxed_rank_002_alphafold2_multimer_v3_model_1_seed_000.pdb 79.41 5.69 |
| b55_Q9Y597-KCTD3_HUMAN_unrelaxed_rank_001_alphafold2_multimer_v3_model_2_seed_000.pdb 65.54 18.64 |
| b55_Q9Y618-NCOR2_HUMAN_unrelaxed_alphafold2_multimer_v3_model_1_seed_000.pdb 66.88 6.69 |
| b55_Q9Y6Q9-NCOA3_HUMAN_unrelaxed_rank_001_alphafold2_multimer_v3_model_4_seed_000.pdb 64.93 6.14 |
